# MemPPI platform for measuring and engineering membrane protein-protein interactions in mammalian cells via split nanoluciferase

**DOI:** 10.1101/2025.06.10.658938

**Authors:** Weiyi Tang, Sebastian Jojoa-Cruz, Jiayi Li, Marco Mravic

## Abstract

Membrane protein–protein interactions (memPPIs) underlie critical biological processes including mediating signal transduction, transport, cell communication, and membrane organization. Despite their importance, systematic and quantitative measurement of protein transmembrane (TM) domains laterally interacting in mammalian cells remains challenging. Conventional fusion protein assays directed at memPPIs suffer from low dynamic range, due to the crowded membrane environment and interference from overexpression, trafficking, or self-complementation artifacts. Here, we assessed and further developed the split luciferase complementation NanoBit system to establish suitable expression constructs and working conditions for robust quantification of TM PPIs within human cells. We benchmarked the platform using a panel of natural and synthetic memPPI pairs with diverse architectures, insertion topologies, and trafficking patterns, finding assay settings that robustly distinguish true positive interactions from co-expressed non-interacting pairs. Adding a novel procedure measuring and normalizing protein expression levels in situ in parallel, we drastically improve assay dynamic range and consistency in discerning relative interaction propensities across broad expression levels – which also enhanced compatibility with protein engineering screens. To extend measurements to versatile protein contexts, we developed membrane expression constructs and validated tagging strategies to improve trafficking with defined TM orientations. By incorporating topological control and accounting for variations in expression levels, this memPPI platform becomes a viable approach for high-throughput interaction analyses and experimental screens for membrane protein design such as TM binders targeting human membrane proteins. Owing to its sensitivity, scalability, and adaptability, we expect this assay platform to have broad cross-disciplinary utility in protein science, design, and drug discovery.

## Introduction

Membrane proteins assemble into functionally important complexes vital for diverse biological processes and regulation. Interactions between proteins within lipid bilayers, whether transient or constitutive, can also be chemically targeted towards therapeutic modulation^1^. Several assay systems exist for measuring membrane-embedded protein-protein interactions (PPI), varying drastically in their dynamic range and adaptability^2,3^. Here, we assess an emerging assay system, NanoBit^4^, for the high-throughput analysis of transmembrane (TM) PPI in mammalian cells focused on those complexes mediated principally by TM regions.

Split protein and 2-hybrid systems fused to TM domains have been extensively explored to characterize homo- or hetero-oligomeric complexes of both natural and engineered assemblies^2^. *E. coli*-based systems like ToxR use TM helices’ ability to self-interact and induce homo-dimerization of a fused transcription factor to initiate genetic reporters for mutually parallel single- and multi-pass membrane proteins^5–8^. Similar colorimetric reporter systems have been modified or developed to measure hetero-dimerization of distinct TM spans, i.e. dominant-negative (dn) ToxR^9^, dnARAC^10^, BACTH^11^, or the LexA-LexA* GALLEX^12^ systems. By instead incorporating an antibiotic resistance reporter, the analogous ToxCAT^13^ assay enables genetic selections of TM span self-assembly in pooled library-based screens^14,15^. As well, a split beta-lactamase (BlaTM) strategy has been used to extend measurement beyond mutually parallel TM interactions to antiparallel complexes while also restricting interaction of consequence to those with periplasmic localized enzyme fragments in the intended insertion topology^16,17^. This suite of assays has been highly effective for studying TM domain interactions, particularly for single-span domains, leading to foundational discoveries concerning the sequence determinants directing interactions in membranes^5,11,13–16,18,19^. However, utilizing *E. coli* as host drastically restricts investigations into most multi-spanning membrane proteins involved in human health and disease.

To test mammalian proteins in eukaryotic cell lines, split protein complementation using enzymes (DHFR^20^, β-galactosidase^21^, firefly luciferase^22^, etc.), fluorescence proteins^23,24^, and others^25–27^ have been used extensively^28^; however, many have drawbacks which are further exacerbated when examining membrane proteins. The decisive factor and typical shortcoming of most split proteins are their level of complementation independent of fusion partner PPI due to inherent self-interaction propensity^29^ and/or irreversibility^30^ that establish the basal signal level and usable dynamic range. Often the signal readout (fluorescence, enzyme activity, antibiotic resistance, metabolite dependence, etc.) over background is not substantial and, when combined with variability in protein expression levels, leads to difficulty in discerning true positive PPIs. The crowded cellular membrane environment can further reduce the signal-to-noise between background collisions and true PPIs, making discerning transmembrane PPIs in mammalian cells using biomolecular fluorescence complementation (BiFC) very difficult^31,32^. Modern assays strive for selectivity by requiring multiple molecular events in series as added layers of control: BRET^33,34^, TANGO^35^, SPARK^36^, mammalian-membrane two-hybrid split ubiquitin complementation (MaMTH) system^37^. This class of multi-step assays are reliably employed for GPCR recruitment of beta-arrestins or G-proteins and inter-cell surface interactions^38^, but are not widely employed for measuring PPIs between two transmembrane proteins. These assay formats suffer from high background when both protein components are membrane-spanning. Likewise, strategies fusing multiple domains to large membrane protein constructs often exhibit shortcomings in trafficking and lack versatility for measurements in alternative contexts (e.g. distinct sub-cellular localizations, insertion topologies, engineered sequences).

We sought to develop an assay to evaluate versatile membrane protein complexes at scale, enabling chemical biology campaigns such as interaction network mapping, drug discovery, or computational protein design. Such measurements create opportunity to connect design principles, atomic-detail rationale, and algorithms for evaluating PPIs in cells – especially for complexes mediated by TM spans. We propose that the split nanoLuciferase (nanoLuc) NanoBit system’s unique characteristics are uniquely positioned for discerning PPI measurements between mutual membrane-spanning proteins partners of diverse topologies. LgBit (19 kDa) and smBit (1.3 kDa) components have minimal self-complementation affinity (K_d_ ∼ mM without interacting fusion partners)^4^, reversible nature, minimal fusion mass, and enzymatic signal amplification – behaviors leading to the observed higher signal-to-noise for most soluble protein than classic PPI assays. Thus far, NanoBit’s performance and technical limitations in such TM-focused application have not yet been thoroughly assessed. Early precedent for monitoring PPIs by NanoBit in the membrane include interactions between membrane proteins^39–41^ and G-proteins recruitment to GPCRs^42^. NanoBit of constitutive GPCR dimers CB_1_R and 5-HT_2A_R demonstrates orders of magnitude better fold-change over background compared to analogous BiFC measurements^32^.

Here, we evaluated the utility, scope, and conditions for NanoBit to reliably distinguish true interactions in the same cell between two membrane proteins from non-interacting co-expressed protein pairs across distinct protein topologies and architectures. Novel constructs for robust trafficking and TM protein engineering in HEK293T cells are presented that control topology, hosting diverse synthetic sequences and appropriately orienting TM spans. Normalization of PPI signal by a relative protein’s expression level was also implemented to better discern interaction propensities, aiding high-throughput screening procedures. Establishing successful, reliable working principles for NanoBit’s application to membrane protein-protein interactions (memPPI) broadens the capabilities to investigate transmembrane interactions and facilitates protein engineering targeting mammalian membrane proteins, e.g. by computational design.

## RESULTS

### MemPPI can distinguish natural and designed membrane protein interactions of distinct topologies

We first sought to establish what is the expected signal-to-noise for a range of relevant true positive natural and synthetically optimized membrane protein complexes, as well as true negative constructs to establish the critical non-interacting baseline from non-specific complementation. Two key determinants in our design were developing constructs where fused LgBit and smBit domains are properly trafficked, localized to the same cellular compartment, and quantitatively monitoring apparent protein expression levels – which depend both on the amount of DNA transfected and each protein’s intrinsic expression and trafficking efficacy. A few isolated studies have monitored NanoBit activity upon transiently transfecting expected TM interaction partners (GPCR dimers^32^, VEGFR2/NRP1^40^, HIV VPU / human CD317^39^, IL-23R dimers^41^). These cases use a relatively high amount of DNA transfected (40-200 ng for each component) ranging on a case-by-case basis from investigator experience. Likewise, positive control PPI levels upon co-expression were typically compared to that of expressing separate components alone (smBit or LgBit) or co-expression of the target membrane protein with a non-membrane protein partner (HALO-smBit or smBit peptide). In test cases for membrane-embedded proteins, the pertinent baseline comparison of nanoLuc complementation upon co-expression of non-interacting proteins relative to a true PPI have not been thoroughly evaluated.

To establish the benchmark for signal-to-noise fold increase for putative positive TM PPIs relative to non-interacting partner pairs, we measured split nanoLuc complementation with two natural complexes of GPCRs with TM accessory proteins of distinct topologies of known structure (**Fig. 1a**): class B calcitonin-like receptor (CLR) and its type I accessory adaptor <u>r</u>eceptor <u>a</u>ctivity-<u>m</u>odifying <u>p</u>rotein 1 (RAMP1)^43^; class A sphingosine 1-phosphate receptor 1 (S1PR1) and its type II adaptor CD69^44^. Likewise, we evaluated two known stable *de novo* designed membrane protein assemblies of single-pass subunits previously untested in the mammalian cell environment (**Fig. 1a**): the type I parallel C5-symmetric homo-pentamer “eVgL”^45,46^ and antiparallel complex between type I mouse erythropoietin receptor (EpoR) and design type II “CHAMP” inhibitor^47^. In the assay, one protein is deemed the “Target”, typically the natural membrane protein, tagged with the minimal smBit fragment at its C-terminus to be minimally perturbative to avoid modifying N-terminal signal peptides. The interaction partner, referred to as the “Binder”, is LgBit-tagged at its intracellular termini and Flag-tagged at the extracellular termini – envisioned to be the construct varied during potential protein engineering screens. We chose a series of mock “Binder” TM proteins known to express well and traffic with high fidelity to the plasma membrane surface to serve as non-interacting negative control to reveal the baseline level of non-specific split enzyme complementation simply from co-expression with a given “Target”: natural CD4 or CD8a, and a *de novo* 3-pass protein. Additionally, positive control “Binders” (RAMP1, eVgL) for one “Target” can be re-used as negative controls for a different “Target” when there is no expected interaction.

**Figure 1.**
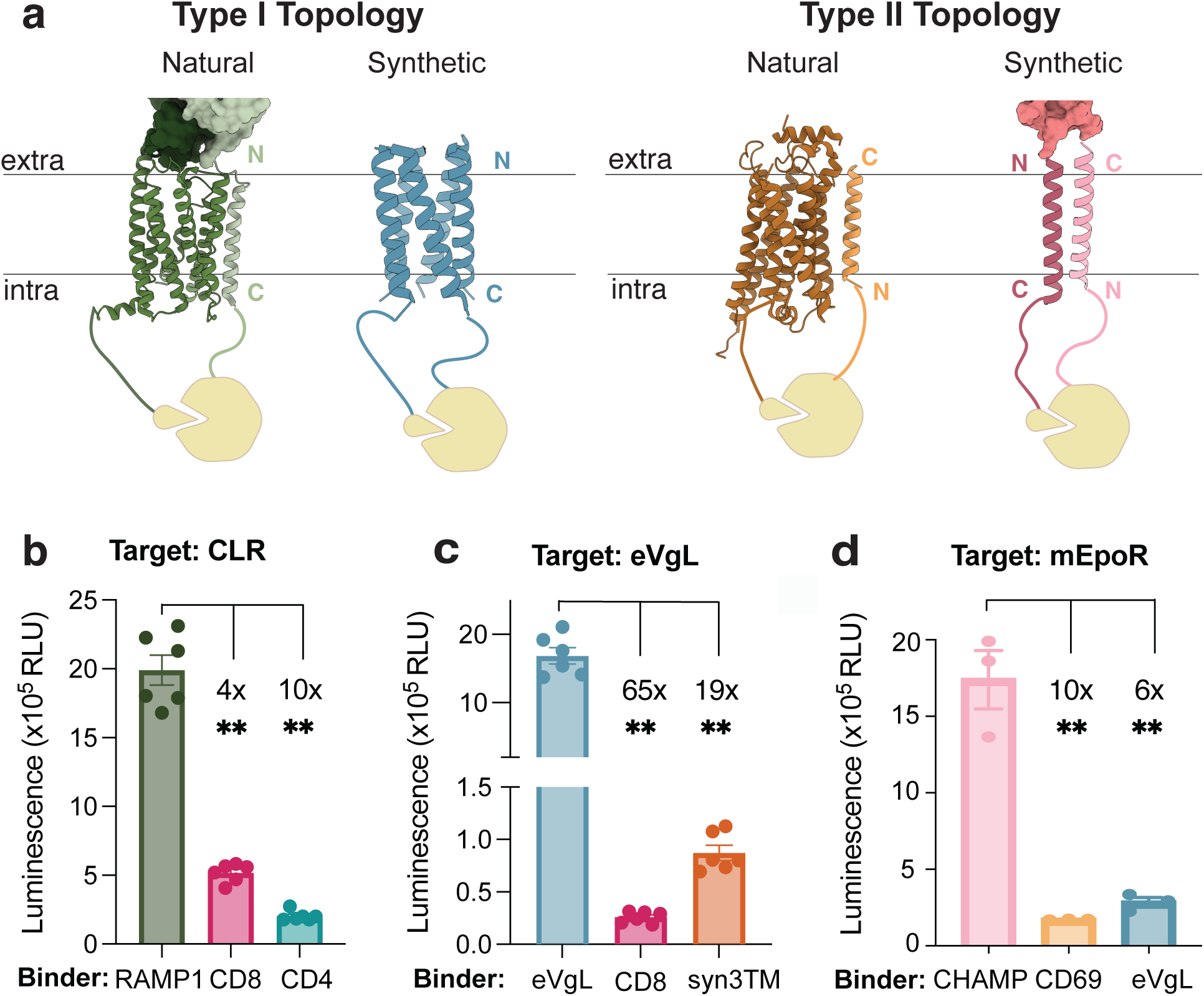
NanoBit enables diverse membrane protein-protein interactions (memPPI) measurements. **a.** Schematic of membrane PPI by NanoBit. For type I binders, CLR (dark green) interacting with its natural binder RAMP1 (light green) (PDB:6E3Y), fused with smBit and LgBit respectively and eVgL, a design protein pentamer, fused with smBit and LgBit (PDB: 6MQU). For type II binders, S1PR1 interacting with CD69 (PDB: 8G94) and EpoR interacting with type II design protein CHAMP. Intra: Intracellular; Extra: Extracellular. N and C denote the N and C termini of the protein. **b.** NanoBit for natural membrane PPI. Luminescence signals with known binder (RAMP1) and non-specific interaction binders (CD8, CD4 and EpoR). Data show n=2 trials, representative N=3 total independent experiments. Error bars, standard error of the mean (SEM). Asterisks indicate P values reaching <0.001 from unpaired t-test compared with CLR-RAMP interaction. **c.** NanoBit for synthetic membrane PPI. eVgL in its homopentamer and with non-specific binders(CD8 and designed protein syn-3TM). n=2 trials shown, representative of N=3 independent experiments; error bar SEM; P value <0.001 (unpaired t-test) compared to eVgL-eVgL interaction. **d.** NanoBit for type II membrane PPI. EpoR with its designed binder CHAMP, and with non-specific binders (CD69 and designed protein eVgL). Data show n=1 trial, representative N=2 total independent experiments; error bar SEM; P value <0.001 (Tukey’s test) compared to EpoR-CHAMP interaction.

In designing optimal NanoBit conditions for signal-to-noise, we posited that only a few copies of each membrane protein would be necessary to detect a protein complex. Conversely, transfecting large amounts of DNA leading to overexpression likely is detrimental to signal-to-noise. For the diverse PPI pairs described, co-expression from 10-30 ng DNA in 96-well plates resulted in strong NanoBit signal for the expected positive PPI pairs (∼10^6^ RLU, **Fig. 1b-d**, **Fig. S1**). Relative to the baseline complementation set for each “Target” by co-expressing non-specific TM protein controls, the fold change increase in NanoBit luminescence spanned 3-10x for known natural complexes and was 60-fold increase for the designed eVgL (p-values < 0.05). By contrast, when large amounts of plasmid are transfected (range of 100-200 ng range per component, typical for lipofectamine and reported for GPCR dimers^32^), we found a drastic impairment in discrimination of known positive PPIs from non-interacting protein pairs (**Fig. S2a**). Co-transfecting 200 ng of “Binder” DNA alongside 100 ng CLR “Target” DNA results in true negative pair CLR–CD8a having greater assay signal than true positive CLR–RAMP1 pair (**Fig. S3b**), only 1.2-fold difference at 100 ng “Binder”, and 2-fold change at 50 ng “Binder”. These data, along with our lab’s broader experience with other protein systems, indicate that NanoBit does not readily work to discern complexes between membrane proteins under standard assay conditions without optimization.

We further tested the hypothesis that lower TM protein expression level might increase the ability to distinguish true positive from non-specific membrane-spanning PPIs through systematic titration of DNA transfected, comparing signal of true positive PPIs eVgL–eVgL and CLR– RAMP1 with that of “Targets” instead co-transfected with non-interacting TM proteins as mock “Binders” (**Fig 2**). For a fixed amount smBit-tagged “Target” eVgL DNA, adding less “Binder” DNA (1:1, 2.5:1, 5:1, 25:1 target:binder mass ratio) resulted in consistently high fold-change between cells with LgBit-tagged eVgL (true positive) compared to non-interacting CD8a (34-80x, n=3, p<0.001). Thus, eVgL self-interaction is clearly discriminated from its random collisions with CD8a and homopentamerization is robust even at low copy numbers (**Fig. 2a**). For CLR, its NanoBit signal with RAMP1 relative to that with non-interacting partners (CD8a, CD4, eVgL) increased as the amount of DNA transfected decreased **(Fig. 2b)**. Fold-change from the non-interacting baseline set by CD8a was modest at 20 ng, 2.16 ± 0.04, (equal Target:Binder DNA). For lower DNA delivered, 10 and 5 ng, fold change was similar, 4-5x, but increased to 13-fold (± 1.45) above baseline for the lowest DNA amount as just 1 ng of “Binder” per well (n=2, p<0.001 for all) (**Fig. 2b**). These results were consistent whether transfection was conducted with empty vectors added as vehicle to equate total DNA transfected or in conditions where “Binder” or “Target” DNA delivered are uneven (**Fig. S3)**. These data further support that keeping membrane protein expression levels low can be beneficial in improving the signal-to-noise observed for TM PPIs.

**Figure 2.**
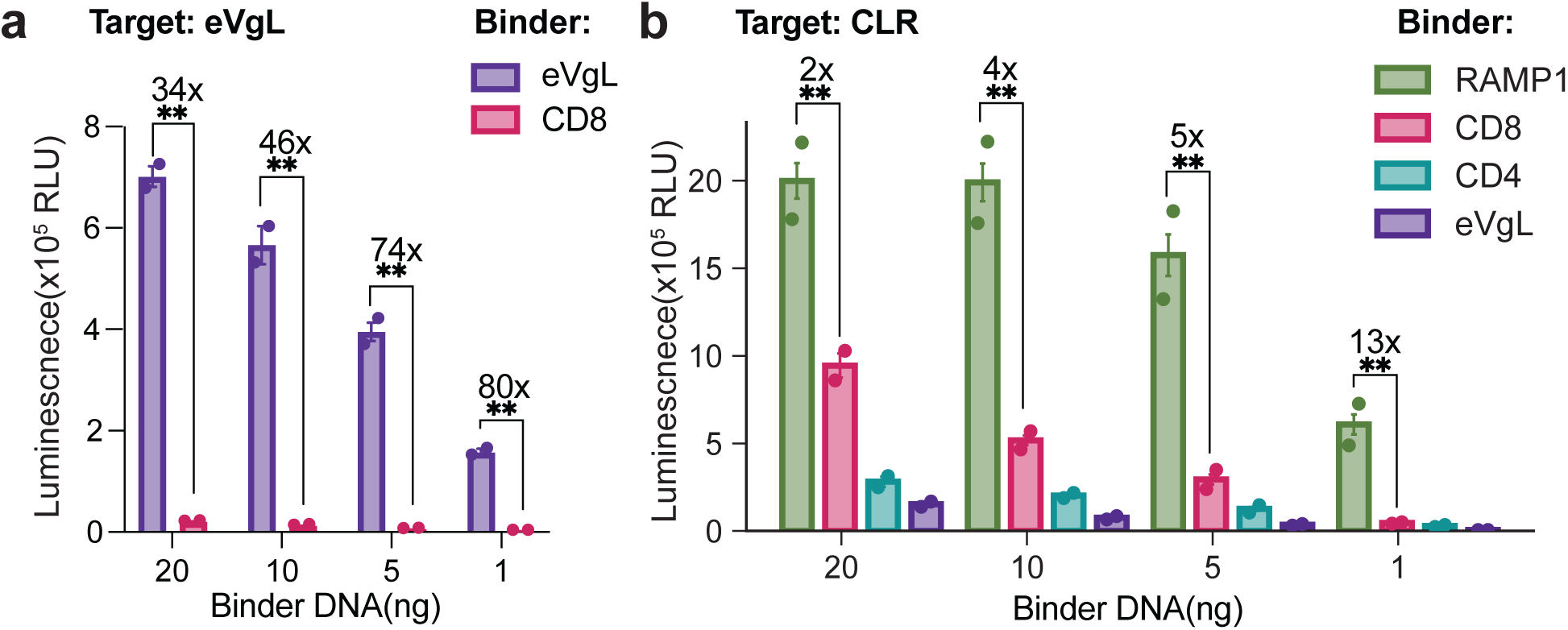
memPPI signal-to-noise range is highly sensitive to transmembrane protein expression level. **a.** eVgL-smBit target (20ng DNA) transiently co-transfected with either eVgL-LgBit and CD8-LgBit binder at varying amount of binder DNA modulating binder protein expression level (20, 10, 5, and 1ng) in HEK293T cells. Raw luminescence signals with fold-change between positive and negative controls noted. Data points are averages of technical triplicates for N=2 independent experiments. Error bars, SEM; Asterisks indicate P values reaching <0.001 from unpaired Welch’s *t*-test. **b.** CLR-smBit target (20ng DNA) co-transfected with natural and synthetic TM LgBit-tagged potential binders upon binder DNA titration (20, 10, 5 and 1ng transfected DNA) in HEK293T cells. Raw Luminescence signals, with fold-change between positive and negative controls noted. Data points are averages of technical triplicates for N=2 independent experiments. Error bars, SEM; P<0.001 from unpaired Welch’s *t*-test.

### Quantitative normalization by binder expression increases consistency and discriminatory power across scales

Given the apparent relationship with expression level and dynamic range of NanoBit between membrane proteins, we devised a strategy for quantification of protein copy number *in situ* to better distinguish whether high luciferase complementation signal occurs simply from higher relative component protein expression levels or from a true PPI. We leveraged the LgBit domain to quantify the “Binder” component, measuring luminescence upon digitonin-facilitated intracellular delivery of the HiBiT peptide. HiBiT’s nanomolar affinity for LgBit, reconstituting nanoLuc spontaneously, should outcompete any expressed smBit. This procedure avoids lysis procedures, which often aggregate membrane proteins, and should report on the total content of accessible LgBit-tagged “Binder” protein in live cells, not just those participating in specific or non-specific PPIs. In the 2-step protocol (**Fig 3a**), first NanoBit luminescence is measured to assess memPPI; second, the HiBiT peptide is introduced, and luminescence is remeasured in the same assay well to quantify total “Binder-LgBit” expression. This approach is easy to implement and especially useful for monitoring very low protein expression levels (recommended for memPPI) near or below detections limits of conventional microscopy and immunoblotting.

**Figure 3.**
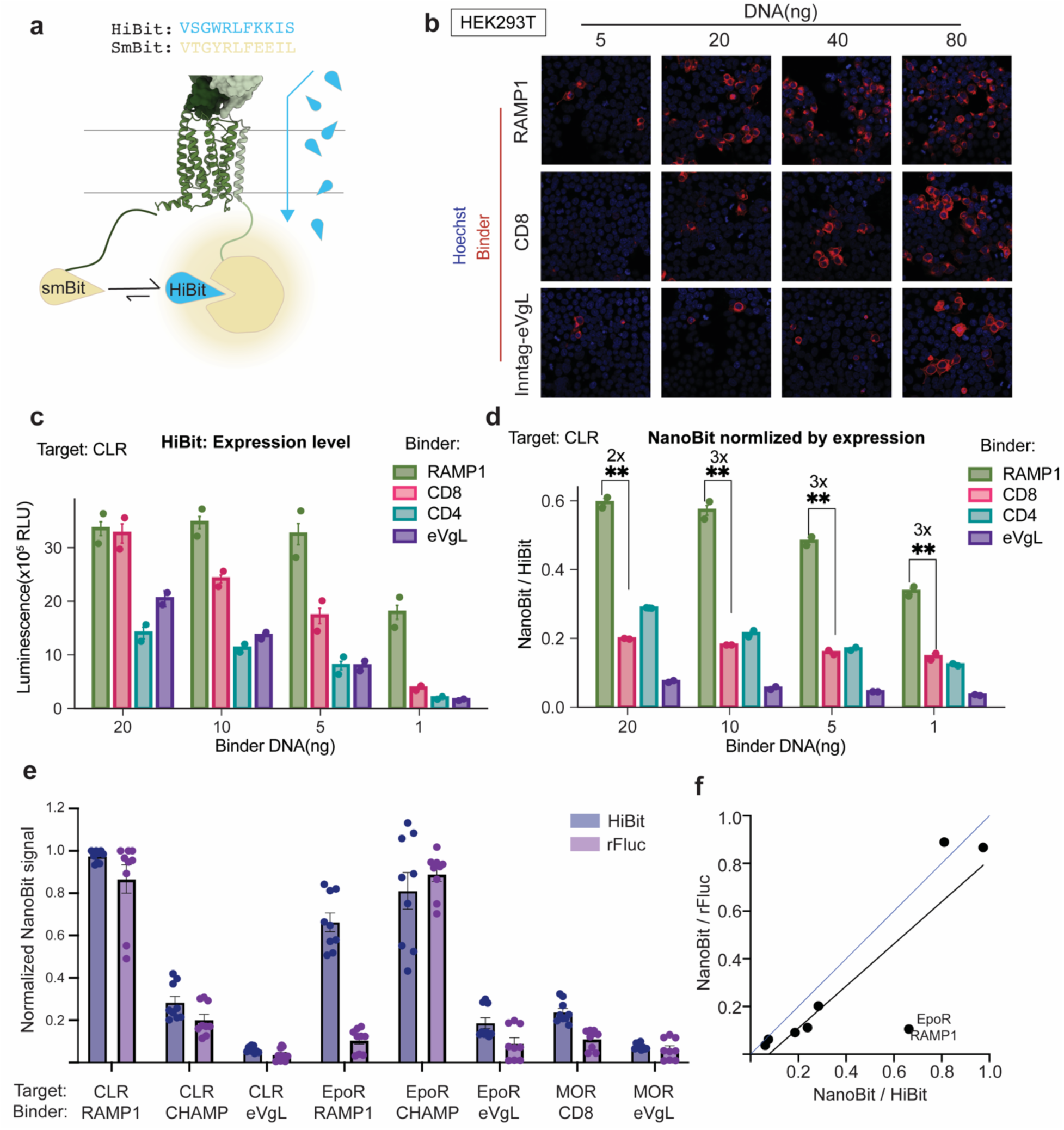
Relative protein expression level normalization via HiBit addition. **a.** Schematic of HiBit peptide (blue) added to cells with permeabilization allows for total LgBit-protein quantification, by its saturation of the LgBit present, out-competing TM protein-tagged smBit (beige). **b.** Immunofluorescence of different LgBit-tagged binders, RAMP1, CD8 and InnTag5-eVgL varying DNA transfection levels (5, 20 ,40, and 80ng), co-expressed with CLR in HEK293T cells, slightly scaled up relative to 96-well plate memPPI assay. **c.** Raw luminescence of LgBit-tagged binders, RAMP1, CD8, CD4, and InnTag5-eVgL upon DNA titration (1, 5, 10 and 20ng) in co-transfection with CLR-smBit upon 1 μM HiBit treatment with 0.001% digitonin semi-permeabilization. Data points are technical triplicate averages from N=2 independent experiments. Error bars, standard error of the mean (SEM). **d.** NanoBit signal normalized by binder expression level (NanoBit / LgBit+HiBit) from CLR-smBit co-transfected with LgBit-tagged binders and binder DNA titration (1, 5, 10 and 20ng). Data points are technical triplicate averages from N=2 independent experiments. Error bars, SEM. **e.** Comparison of NanoBit assay normalized by LgBit-tagged protein expression level measured either directly by HiBit-treated nanoLuc complementation(LgBit+HiBit) or indirect quantification by rFluc expressed downstream of the binder (T2A-rFLuc). Several target-binder pairs measured N=3 independent trials of technical triplicates. **f.** Correlation of mean nanoBit signal normalized by direct HiBit addition or indirect T2A-rFluc quantification of samples in panel **e**. Black line, linear regression line with R^2^=0.74 and Pearson correlation coefficient of 0.86; blue line, x=y regression line representing an ideal correlation. EpoR–RAMP1 pair is an outlier, caused by higher rFluc luminescence relative to HiBit detection.

To benchmark adding this LgBit-HiBit quantification step to the NanoBit assay, we first evaluated the expression levels by immunofluorescence microscopy for 3 example “Binder” constructs co-expressed with “Target” CLR in transfection conditions proportional to 96-well NanoBit: RAMP1, CD8a, and eVgL fused at the N-terminus with the InnTag5 domain (InnTag5-eVgL improved plasma membrane trafficking versus untagged eVgL from Figures 1-2). CD8a exhibits a somewhat linear trend in expression level as more DNA is transfected, whereas RAMP1 plateaus at high expression early in the DNA titration (**Fig 3b**). InnTag5-eVgL expression remains low until the highest DNA level. When monitoring LgBit luminescence in HiBit-treated cells under analogous conditions co-expressing various amounts of “Binders” with CLR, the signal (indicative of expression level) relative to DNA transfected mirrors each construct’s behavior observed by microscopy (**Fig 3c**). We then integrated this HiBit-LgBit measurement to normalize the preceding memPPI NanoBit signal, resulting in an interaction propensity relative to the amount of LgBit-Binder expressed. Traditional NanoBit comparing CLR–RAMP1 and CLR–CD8a co-expressed pairs yielded differences in luminescence depending heavily upon the nanogram (ng) of DNA delivered, ranging between 2 to 13-fold (**Fig 2b, 3c**). After normalizing by the HiBit-LgBit expression level, the ratio of normalized PPI signals for CLR–RAMP1 versus negative PPI pairs becomes far more consistent across DNA transfection conditions **(Fig. 3d)**: e.g., CLR–RAMP1 versus CLR–CD8a, 2.1 ± 0.1, 2.7 ± 0.1, 2.9 ± 0.1, 2.7 ± 0.2 fold-changes for 20, 10, 5, and 1 ng Binder, respectively; all p < 0.001; n=2).

To further evaluate the normalization of NanoBit signal by protein expression level, we tested the HiBit-LgBit method for 8 additional PPI pairs and in comparison with a canonical co-translational reporter approach indirectly tracking an orthogonal rFluc downstream of the “Binder” (“Binder”-T2A-rFluc). That includes 2 known positives (CLR–RAMP1, EpoR–CHAMP) and 6 expected negatives with targets CLR, mu-opioid receptor (MOR), and EpoR (**Fig 3e**). The NanoBit luminescence, whether normalized by rFluc or HiBit-LgBit mechanisms, are in strong agreement for both positive and negative PPIs (overall Pearson correlation coefficient = 0.86; **Fig 3f**). Thus, similar performance for discriminating positive PPIs is achieved, with consistent relative “Binder” expression measurements for 7/8 PPI pairs (**Fig S4**). The outlier PPI, EpoR–RAMP1, has not been previously detected or otherwise measured to our knowledge. This Target-Binder pair exhibits modest NanoBit PPI levels, but when normalized by rFluc appears as low (negative PPI) or as high by HiBit-LgBit (positive PPI) (**Fig S4**). It is unclear whether this PPI is a false negative from rFluc detection, false positive from HiBit-LgBit, or a new true PPI, but it is noted that RAMPs are chaperones with broad scopes of interaction, immunoprecipitating with several receptor classes^48,49^. Nonetheless, the data supports utility of 2-step NanoBit method for normalizing by expression level. The HiBit-LgBit method holds advantages in the context of membrane protein trafficking, avoiding 2A peptides, and in that direct *in situ* measurements should be more accurate for dynamic processes and scenarios where co-transcriptional or co-translational reporters mismatch endpoint protein levels. This simple method executed on the same cells (same assay well) reading out LgBit-fused protein copy number on the fly, addresses the challenge of comparing absolute NanoBit signals and interpreting PPIs across different pairs which inevitably have variance in underlying proteins’ expression levels.

### Engineering membrane protein topological control for relevant PPI measurements

An important underlying assumption for relevant measurements is that the membrane proteins are expressed (detectable by LgBit/HiBit), trafficked correctly, and topologically oriented to correctly position the LgBit domain in the same cellular compartment as the smBit tag. To facilitate this behavior, membrane protein constructs must be evaluated and often improved by optimal sequences of signal peptides, epitope sites, soluble domains, and other tags. These are difficult facets to control at protein engineering scales, screening many different TM architectures or pools of diverse (possibly *de novo*) sequences. We report 3 practical case studies emphasizing the need for suitable protein constructs and how working principles of membrane protein trafficking can be incorporated to make assaying several natural and synthetic transmembrane domains accessible by memPPI.

First, we sought to develop robust protein scaffold constructs able to host diverse natural or synthetic TM sequences which direct Type II protein insertion: reliably trafficking those TM spans into an orientation antiparallel relative to natural Type I proteins like cytokine and immune receptors. We developed a series of constructs to enable measurements screening putative synthetic receptor complexes, benchmarked using the recent protein design success case of TM protein CHAMP and its antiparallel-oriented complex with EpoR, which inhibits receptor signaling^47^.

Construct design was divided into three main components: soluble domains, N-terminal signal peptide, and TM-proximal linkers. Type II membrane proteins lack a cleavable N-terminal signal peptide, instead having modest-sized polar leader sequence facilitating their first TM segment being post-translationally inverted i.e. N-terminus facing the cytosol. Recent work suggests a sufficiently long C-terminus is favorable for this energetic transition^50,51^. Accordingly, we performed an informatics mining of UniProt database for type II single-pass membrane proteins (SL-9906) mining for plasma membrane localization, ≥3 predicted glycosylation sites supporting that localizing, and those with ≥110 residues of C-terminal region after its TM domain also expected to adopt a non-catalytic globular domain (avoiding undesired activities). After filtering out constitutive oligomers, we identified two candidates as C-terminal extracellular regions for our constructs, human C-type lectin domain family 2 member B (HsCLC2B) and natural killer cell receptor E (HsNKG2-E). We included the monomeric InnTag5, validated for minimal interference protein tagging^52^, and LgBit as additional candidate extracellular domains. For the intracellular N-terminal leader sequences and domains, we varied from a minimal polar segment with an ALFA tag (no folded domain) to including the LgBit domain for assay readout. We selected one short and one long N-terminal leader sequence from our bioinformatic search to facilitate type II protein topology: “ssPolar1” from HsNKG2-E and “ssPolar2” Oxidized low-density lipoprotein receptor 1 (HsOLR1)^53^ based on strength of evidence of type II orientation. Inter-domain linkers were designed to be flexible: the intracellular linker following the positive-inside rule and the extracellular linker having net negative charge. A control construct using the hemagglutinin signal sequence (ssHA) was included, anticipated to yield type I insertion. Three type II constructs were designed which were predicted *in silico* with strong confidence to adopt the desired topology^54^ and localization^55^: positioning the aforementioned modules including LgBit either in the intracellular or extracellular space and a FLAG-tag on the expected extracellular terminus (**Fig 4a**, **Table S4**). Localization of the constructs was tested by immunofluorescence upon transfection into HEK293T cells (polycistronic constructs co-expressing mCherry), staining with or without membrane permeabilization to assess FLAG accessibility and thus construct orientation (**Fig 4b**). The ssHA Construct-1 stains minimally when non-permeabilized, but presents a strong signal for anti-FLAG antibody when permeabilized, indicating intracellular localization i.e. Type I plasma membrane insertion – aligned with expectations. By contrast, Constructs-2 and -3 hosting the ssPolar1 both show surface staining in non-permeabilize cells, indicating a significant amount of extracellular FLAG epitope at the plasma membrane and Type II oriented TM protein as designed. Construct-4 with ssPolar2 showed similar localization pattern as construct 1: minimal extracellular FLAG presentation. Construct-2 shows superior surface expression and trafficking compared with Construct-3, lacking a folded domain at its N-terminus and having extracellular LgBit. However, Construct-3 was chosen for further studies due to intracellular positioning of LgBit and compatibility with intracellular smBit of our existing “Target” constructs.

**Figure 4.**
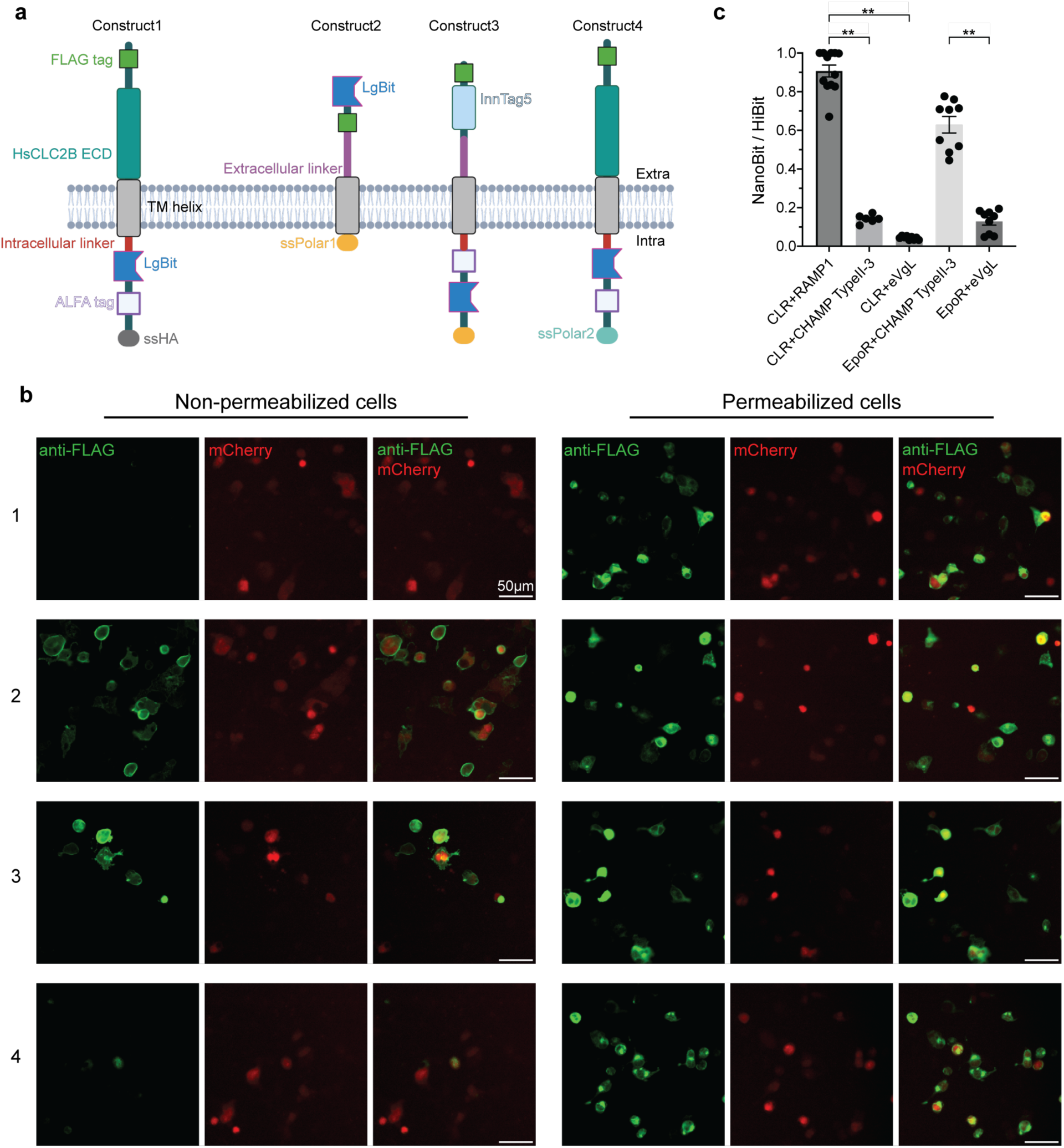
Type II construct design. **a. b.** Schematic and surface staining for type II construct variants of CHAMP. **b.** Normalized NanoBit / HiBit for CHAMP in type II - construct 3 with non-target CLR and target EpoR. Positive control for assay, CLR+RAMP1; negative control for CLR target, eVgL; negative control for EpoR target, eVgL. N=3 total independent experiments with technical triplicates. Data for each independent experiment normalized to its highest value of CLR+RAMP1. Scatters represent each data point across all experiments. Error bars, standard error of the mean (SEM). ** indicates P values <0.001 from Dunnett’s T3 test.

To assess the viability of Construct-3 for memPPI, we measured its ability to host CHAMP to allow interaction with EpoR (**Fig. 4c**). As expected, the EpoR–CHAMP pair resulted in high normalized PPI signal while negative control CLR–CHAMP did not. Thus, we successfully designed and validated Type II single-pass membrane protein constructs compatible with the memPPI assay. Simply having a polar stretch (ssPolar2) the N-terminal leader appears to be insufficient to direct this topology, and the ssPolar1 emerged as a key element determining Type II trafficking. These novel constructs should enable exertion over topological control for variable synthetic or natural TM spans with robust trafficking, with options to position LgBit inside or outside the cell – applicable broadly within protein or cellular engineering applications and for further customization.

### Modulating membrane protein expression, trafficking, and sub-cellular PPI specificity

RAMP1 proteins interact with a broad scope of GPCRs^48^. For the CLR-RAMP1 complex (CGRPR), their TM domains directly interact alongside extensive extracellular regions^43^, whereas the glucagon receptor’s structure with RAMP2 lacked the latter’s TM domain density and suggested extracellular domains mediate their interactions^56^. We sought to use NanoBit to determine if RAMP1 TM span alone is sufficient for interaction with CLR (**Fig. 5a**). However, a construct of the minimal TM span for RAMP1 (C-terminal LgBit) was not adequately expressed (**Fig. 5c**), preventing this test. Alternatively, we hypothesized a globular domain preceding RAMP1’s TM span, akin to its extracellular domain, may aid its correct trafficking and expression. Two RAMP1 chimeras using InnTag5 and HALO as mock extracellular domains both expressed at similar levels as wildtype (WT) RAMP1 with sufficient plasma membrane localization to display the RAMP1 TM span alongside CLR (**Fig. 5a,c**). InnTag5-RAMP1-TM and HALO-RAMP1-TM exhibited 2-fold reduction in NanoBit signal normalized by expression level (HiBit method) relative to the WT CLR–RAMP1 PPI (**Fig. 5e**), being only slightly higher than non-specific interaction pairs CLR–CD8a or CLR–CD4 NanoBit signals (3-fold less than CLR– RAMP1, **Fig. 3d**). Thus, while RAMP1’s TM span may have some weak non-random level of interaction above baseline, the extracellular domain appears to be critical for driving the strong observed complex with CLR. As well, we find addition of innocuous stable soluble domains is a useful strategy to assist in expression of minimal TM domains to facilitate NanoBit measurements and for isolating their contributions to membrane protein interactions.

**Figure 5.**
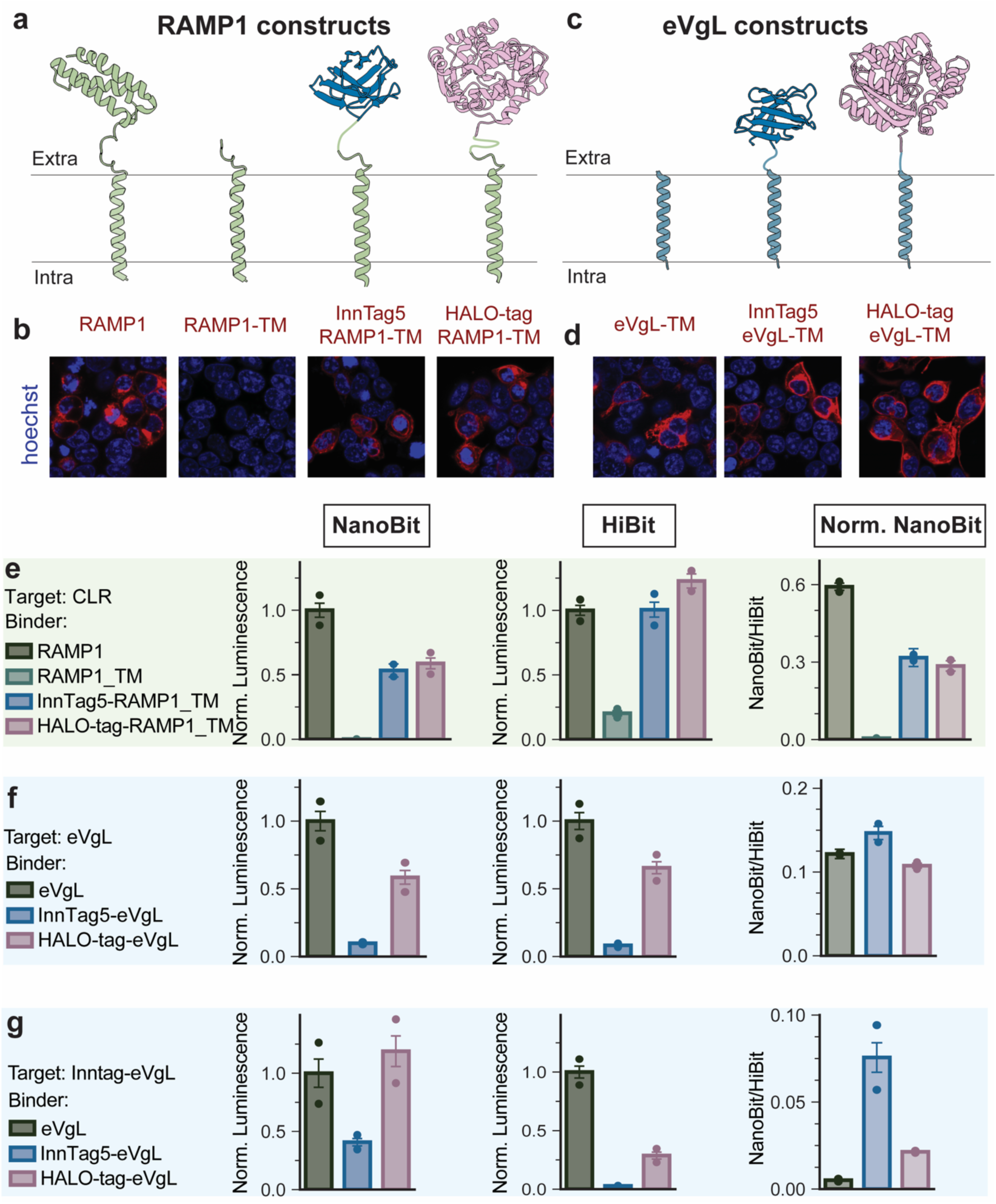
Membrane protein construct trafficking leads to different apparent binding. **a. b.** Schematic and surface staining for construct variants for display of RAMP1, including InnTag5 and HALO-tag variants alongside minimal TM domain variants. Intra: Intracellular; Extra: Extracellular. **c.d** Schematic and surface staining for construct variants for display of synthetic eVgL transmembrane domain including InnTag5 and HALO-tag variants **e.** Luminescence measurement of construct variants for RAMP1 targeting CLR. NanoBit and HiBit signals are normalized to CLR-RAMP. Error bars, SEM; Data representative of N=2 independent experiments; Scatters represent the mean value of each experiment; Norm. Luminescence: Normalized Luminescence. Norm. NanoBit: NanoBit normalized by LgBit+HiBit. **f.** Luminescence measurement of construct variants for eVgL targeting eVgL-smBit. NanoBit and HiBit signals are normalized to eVgL-eVgL. Error bars, SEM; Data representative of N=2 independent experiments; Scatters represent the mean value of each experiment; Norm. Luminescence: Normalized Luminescence; Norm. NanoBit: NanoBit normalized by LgBit+HiBit. **g.** Luminescence measurement of construct variants for eVgL targeting at Inntag-eVgL. NanoBit and HiBit signals are normalized to eVgL-Inntag-eVgL. Error bars, SEM; Data representative of N=2 independent experiments; Scatters represent the mean value of each experiment; Norm. Luminescence: Normalized Luminescence; Norm. NanoBit: NanoBit normalized by LgBit+HiBit.

In testing the *de novo* designed eVgL expressed as a direct LgBit fusion lacking a soluble domain, we find that, despite bearing a ssHA signal peptide, it appears to localize primarily to intracellular membranes rather than the cell surface (**Fig. 5b,d**). Its exceptionally high expression level and presence of intracellular puncta by microscopy cast doubt on whether its high PPI NanoBit signal arises mainly from homo-pentamerization rather than overexpression or intracellular aggregation artifacts. As for RAMP1, we tested fusion of InnTag5 and Halo to eVgL-LgBit-tagged “Binder” constructs for improved plasma membrane trafficking (**Fig. 5b,d**). Both domains drastically increased eVgL’s cell surface localized fraction, although overall expression levels were reduced (**Fig. 5f,g**). With untagged eVgL-smBit as the “Target”, HALO-eVgL and InnTag5-eVgL exhibit similar overall PPI signal by NanoBit after normalizing for protein expression level (HiBit method), indicating the complex is formed somewhere within the cell (**Fig. 5f**). Conversely, when InnTag5-eVgL-smBit is used as the “Target” construct, the substantially surface localized InnTag5-& HALO-eVgL “Binder” constructs with surface localization have largely higher normalized PPI signal compared to the intracellularly localized untagged eVgL-LgBit, whose normalized luminescence appears to be a true negative non-interacting pair (**Fig. 5g**). As with Type II CHAMP–EpoR, studies with eVgL constructs indicate NanoBit exhibits some selectivity for co-localization to the same cell membrane compartment to detect a PPI complex. These case studies exemplify folding considerations and protein engineering strategies for obtaining appropriate constructs with suitable trafficking for natural or synthetic TM interactions, including how to host and display TM spans in mammalian cells when engineering *de novo* protein complexes.

### MemPPI can distinguish specificity between close GPCR homolog complexes

We next tested the applicability of memPPI to detect subtle chemical changes by assaying interaction selectivity between natural protein homologs: the TM adaptor complex of CD69 that favors S1PR1 versus S1PR2 (**Fig. 6a,b**)^44^. Expression normalized NanoBit of CD69 was 2-fold when paired with S1PR1 relative to S1PR2 (**Fig. 6c**, n=3, p<0.001), aligned with previous co-immunoprecipitation experiments finding CD69 interacts more strongly with S1PR1^44^. We then assessed whether memPPI could discern the consequences of CD69 mutations on S1PR1 interaction as previously described^44^. NanoBit of the V48F, V49F CD69 double mutant with S1PR1 showed significant 30% decrease in the normalized signal relative to WT (**Fig. 6c,d**, n=3, p<0.05). Finally, S1PR1-like mutations were introduced to S1PR2 to potentially reconstitute a synthetic interface with CD69^44^. The L157V-L168M variant showed a modest but consistent 15% increase in PPI signal relative to the WT S1PR2’s interaction (**Fig. 6c,e**; n=3, p<0.001). This case study demonstrates that the memPPI assay can measure subtle changes in interaction strength between known PPI and probe selectivity within homologous or mutant protein panels, illustrating broad scope of use from interface mapping of important interface residues and interactome analysis.

**Figure 6.**
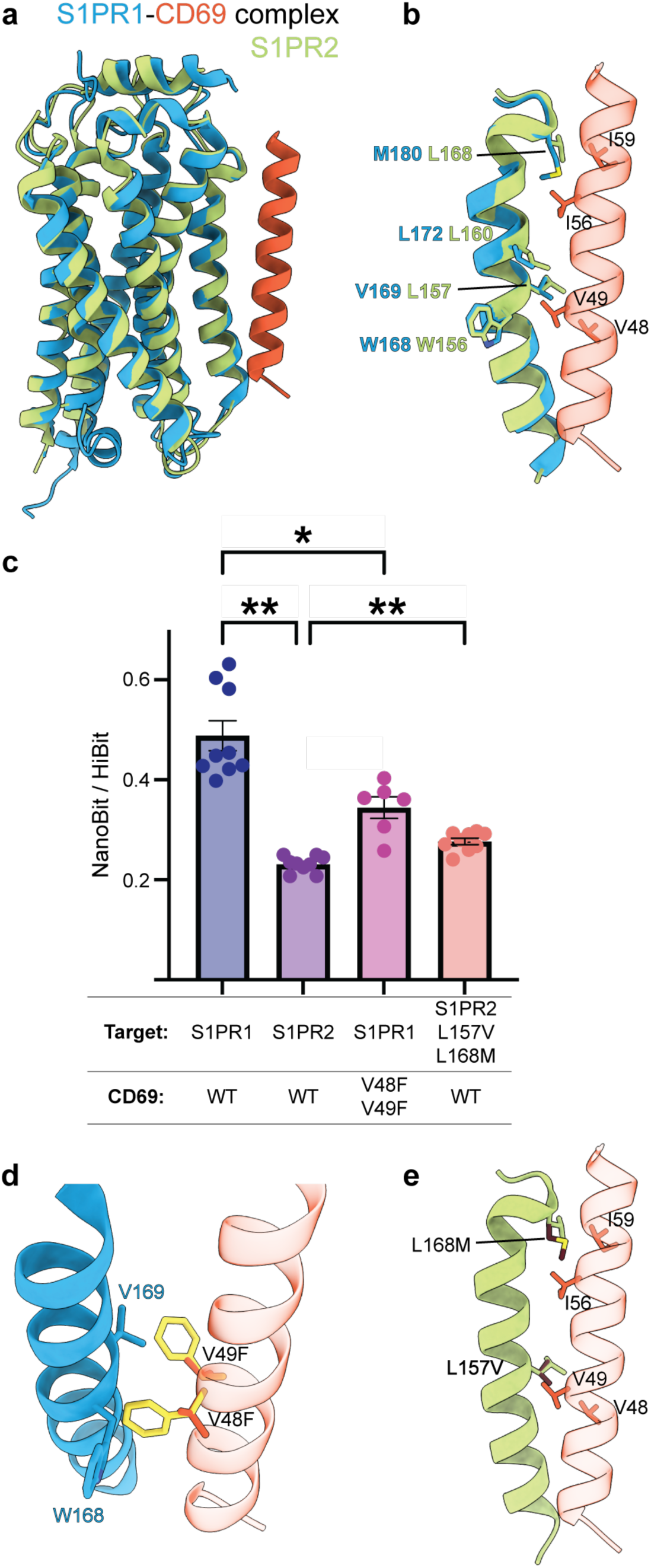
Use of NanoBit / HiBit assay to screen PPI between related targets and effect of mutations. **a.** Structure of S1PR1-CD69 complex (PDB: 8G94) superimposed with S1PR2 receptor (PDB: 7T6B) from the same family. **b.** Highlight of relevant interactions between CD69 and transmembrane helix 4 of S1PR1, and corresponding residues in S1PR2. **c.** NanoBit / HiBit assay results of S1PR1 and S1PR2 with CD69, and effect of selected mutations. N=2 (for S1PR1 + CD69 V48F V49F mutant) or 3 total independent experiments with technical triplicates. Data for each independent experiment normalized to its highest value of CLR+RAMP1 (not shown). Scatters represent each data point across all experiments. Error bars, standard error of the mean (SEM). ** indicates P values < 0.001, and * indicates P < 0.05 (Dunnett’s T3 test). **d.** Model of V48F V49F mutations in CD69 and potential effect on its interaction with S1PR1. **e.** Model of L157V L168M mutations in S1PR2 to resemble S1PR1 and favor its interaction with CD69.

### In Situ Expression Normalization enables scalable high-throughput protein engineering screens

Finally, we leveraged the protein normalization step of our 2-step memPPI and evaluated its performance in an accelerated screening pipeline testing a panel of computationally designed TM domains with intended Type II topology as binders for a GPCR of interest, referred to as GPCR-A. From a pool of designed proteins, cloned in the aforementioned Construct-3 in *E. coli*, individual clones were subject to plasmid amplification by Rolling Circle Amplification (RCA). Unpurified diluted RCA product was used directly to co-transfect library members alongside 5 ng GPCR-A-smBit to HEK293T cells to read the NanoBit PPI alongside protein expression levels (**Fig 7a**). Thus, the laborious step of plasmid purification was bypassed and resulting protein expression level is determined *in situ* in lieu of knowing the precise “Binder” genetic material delivered. RCA products can be sequenced individually or by barcoding and next-generation sequencing (NGS). Interestingly, for the included true positive control PPIs EpoR–CHAMP and CLR–RAMP1, transfection with the RCA product results in the same normalized memPPI signal as delivering approximately 10ng of “Binder” plasmid as input DNA, despite considerable differences in raw luminescence values (**Fig. 7b,c**). Across Target-Binder pairs, although LgBit-HiBit quantification shows order of magnitudes of difference in binder expression levels, the normalized PPI signal compared between plasmid-versus RCA-based transfection is highly similar with excellent linear correlation, R^2^=0.98 (**Fig. 7c,d**). These results establish that normalization by “Binder” protein expression yields self-consistent PPI measurements across an extensive dynamic range and that relative protein levels can be determined *in situ* on the fly in a protein engineering screen. This adaptability facilitates memPPI’s integration into high-throughput platforms and opens opportunities for protein design and engineering screens of synthetic TM domain complexes.

**Figure 7.**
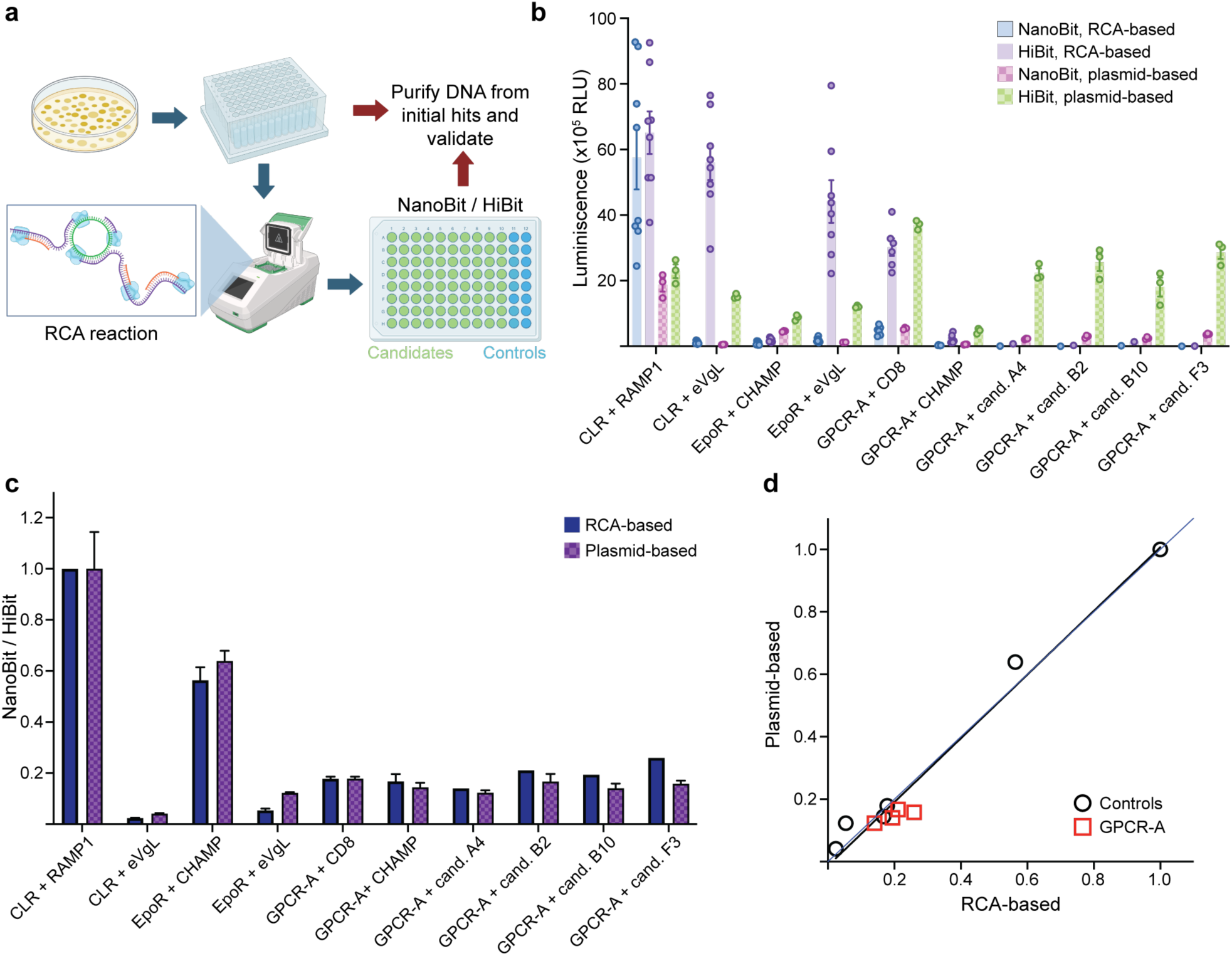
Use of RCA DNA with NanoBit / HiBit assay for fast screening of PPI. **a.** Schematic of pipeline for fast screening of PPI. Colonies transformed with candidate binders of interest are resuspended in a 96-deep well plate, and 1uL of resuspension is denatured and subsequently used as input for the RCA reaction. RCA product is then diluted and used as input DNA of the binder for the NanoBit / HiBit assay. Hits are traced back to the deep well plate, DNA of hits is purified and further validated. **b.** Comparison of luminescence levels of NanoBit and HiBit obtained from using RCA product or plasmid as DNA input for the binder. For experiments with RCA as input, N=2 total independent experiments with technical duplicates for the controls, or N=1 for the tested binder candidates; for experiments with plasmid input, N=3 total independent experiments with technical triplicates. Scatters represent each data point across all experiments. Error bars, standard error of the mean (SEM). **c.** Comparison of NanoBit signal normalized by expression via HiBit for experiments using RCA product or plasmid as DNA input of binder shown in panel **b**. Error bars, standard error of the mean (SEM). **d.** Correlation of mean NanoBit signal normalized by direct HiBit addition for experiments using RCA product or plasmid as DNA input of binder shown in panel **c**. Black line, linear regression line with R^2^=0.97 and Pearson correlation coefficient of 0.98; blue line, x=y regression line representing an ideal correlation.

## Discussion

Here we show that NanoBit can be successfully expanded from soluble or membrane-proximal PPIs to pairs of transmembrane proteins under specific working conditions controlling for common artifacts: over-expression, mis-trafficking, confident non-interacting baseline, and normalizing expression level variability across samples. The motivation to extend and validate NanoBit application scope comes from our challenges faced applying alternative PPI assays to measure TM complex of diverse topologies. This led to establishing a convincing, robust, fast (transient transfection), and scalable platform in well-plate format tailored for memPPIs. To accommodate high-throughput applications, i.e. screening natural interactomes or engineered TM protein pools, we developed expression constructs and tagging strategies promoting reliable, robust protein trafficking, hopefully adaptable to diverse iterations of “Target” or “Binder” proteins. Likewise, developing NanoBit into a 2-step protocol quantifying and normalizing by protein expression levels in the same well dramatically broadens the assay’s practical dynamic range, and reduces technical burden of manually titrating multiple proteins to common expression levels or rigorous quantification of expression in parallel by other means. The resulting measure of relative interaction propensity enables more consistent comparisons even among cell populations varying widely in protein copy numbers. Notably, the confidence to quantify the LgBit-fused component *in situ* regardless of DNA amount delivered or expression efficacy of individual constructs makes our 2-step NanoBit implementation amendable to expedient approaches screening PPIs, including applications designing TM proteins targeting mammalian GPCRs, chaperones, and ion channels.

It is important that users also acknowledge and account for the assay’s current limitations. Just because the smBit-tagged “Target” protein is not similarly directly monitored in the current implementation, it should not be assumed that its expression level and capacity for complementation will be constant when co-expressed with a battery of different TM protein partners. This assumption does holds true for most “Targets” we have tested, but certainly not all. For example, RAMP1 exhibits chaperone capabilities for many GPCRs and increasing expression level and surface localization, including CLR. If the LgBit-tagged component simply alters the smBit component’s levels and basal complementation, this behavior may confound PPI comparisons. Likewise, relative protein stoichiometries, receptor densities, or protein-lipid molar ratios are not quantitatively addressed in this assay format, preventing rigorous assessments of dissociation constants or interaction strength from PPI assay signal – a trade-off for throughput and a native cellular environment. As well, TM protein sequences vary drastically in their characteristic expression, folding and trafficking efficacy under identical transfection conditions (DNA delivered), even for variants hosted in identical construct plasmids. These differences are difficult to rationally control, and are expected to vary across host cell types and when varying TM span sequences – arising from transcriptional, translation, protein folding, stability, and trafficking events. While we validate several constructs and strategies for hosting TM spans, extensive trial-and-error may be necessary to devise constructs with appropriate expression, trafficking, and orientation to assay the wide breadth of natural or synthetic TM topologies of interest. Although tagging with smBit is typically considered to be minimally perturbative, fusion position of LgBit or smBit may disrupt or alter expression patterns – as observed between Type II constructs 2 and 3 (**Fig. 4d**). Likewise, the flexible linkers commonly employed bridging smBit and LgBit domains make complementation events highly permissive and may equally represent direct side-chain directed complexes we intend to assay and broader nanometer-scale co-localization. Thus, the apparent memPPI signal can arise from multiple cellular factors including preferential localization to lipid bilayer microdomains of certain thicknesses or compositions (tetraspanin webs^57^, lipid rafts, cholesterol-rich regions, etc) or from possible shared membrane protein sorting pathways, i.e. prior to reaching the plasma membrane. Thus, all putative complexes should be subsequently interrogated biochemically. While our studies measure a degree of specificity for subcellular localization, NanoBit is a whole cell bulk average measurement, and will report on wherever constructs might co-localize for smBit and LgBit collision – not just at the plasma membrane surface.

Considering these limitations, we propose several practical working principles for assay implementation and interpretation. First, NanoBit experiments should be supported with ample verification of expression levels and trafficking patterns, especially for new “Target” constructs, as done in this study by immunostaining. We evaluate expression levels of a smBit component both alone and under co-expression conditions with a representative set of LgBit-tagged constructs to account for potential co-variation. Although not demonstrated here, T2A-rFluc quantification or flow cytometry may be useful to normalize or collect cell populations to control expression levels of the smBit component for more precise comparisons. Given possible variation in smBit or LgBit levels across protein pairs, we usually recommend testing multiple non-interacting negative control TM proteins be assayed in parallel per “Target”. If possible, having a true positive PPI for that “Target” to assess level of luminescence expected for the high and baseline levels of complementation to compare unknown PPIs pairs is useful. Water-soluble binding domains like nanobodies can be useful positive controls if known TM PPIs are lacking. In cases discovering new PPIs to a “Target” where no true positive control complex exists, we consider a 2-fold expression-normalized luminescence change above the baseline signal of relevant non-interacting pairs with the same “Target” as the conservative minimum indicative of a true complex. Optimized or constitutive high affinity pairs reach into 10’s or 100’s fold-change^32^. For known complexes, smaller but statistically significant decreases in NanoBit signal upon chemical changes may be interpretable as altered interaction relative strength, if expression and localization are equal. We advocate for assay conditions trending towards lower total protein expression levels for increased sensitivity to distinguish true PPIs apart from coincident co-localization or over-expression, although reducing either protein partner below the level of confident expression or detection may lead to false negatives.

## Author contributions

**Weiyi Tang:** Conceptualization; investigation; writing; methodology; visualization; formal analysis; data curation. **Sebastian Jojoa-Cruz:** Conceptualization; investigation; writing; methodology; visualization; formal analysis; data curation. **Jiayi Li:** Investigation. **Marco Mravic:** Conceptualization; writing; methodology; visualization; formal analysis; data curation, supervision.

## Supplementary Material

**Supplementary Figure 1.**
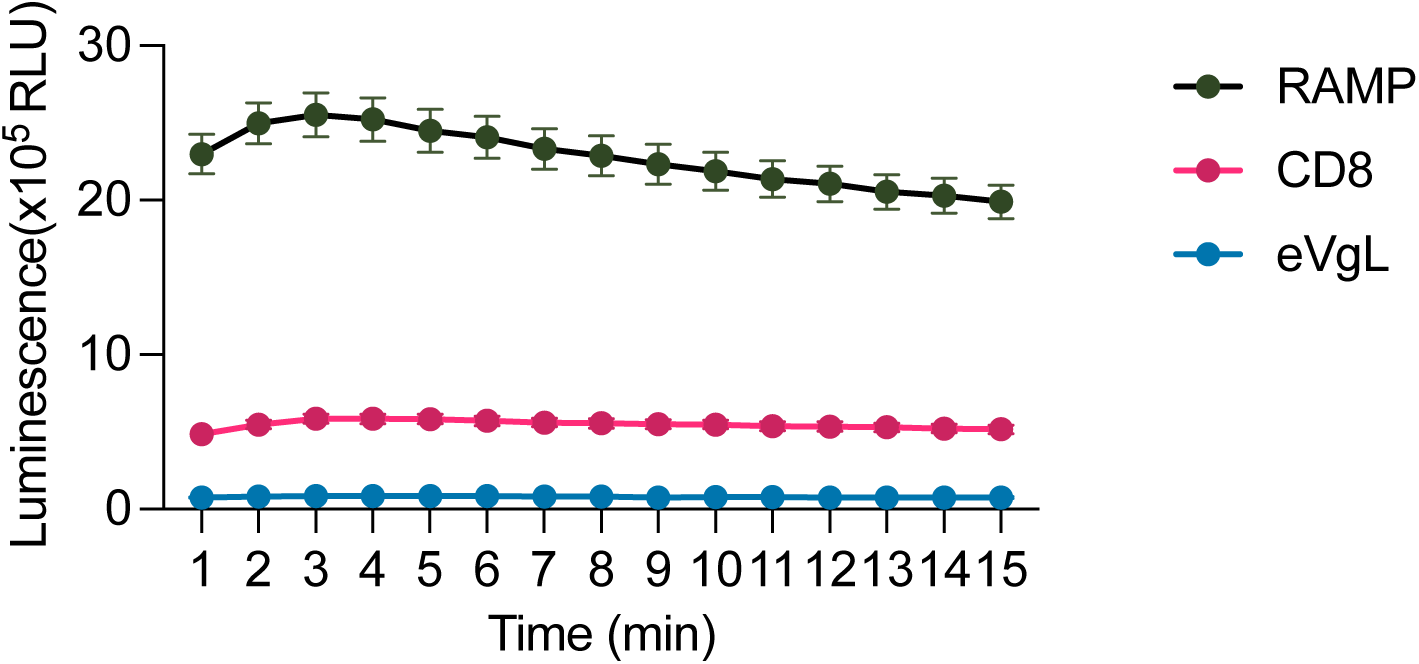
NanoBit glow kinetics. Data represent raw luminescence collected for 15 minutes after exchanging media into 10 µM furimazine in CO_2_-independent media. Data show n=2 biological replicates, representative of n=3 total independent experiments. Error bars, standard error of the mean.

**Supplementary Figure 2.**
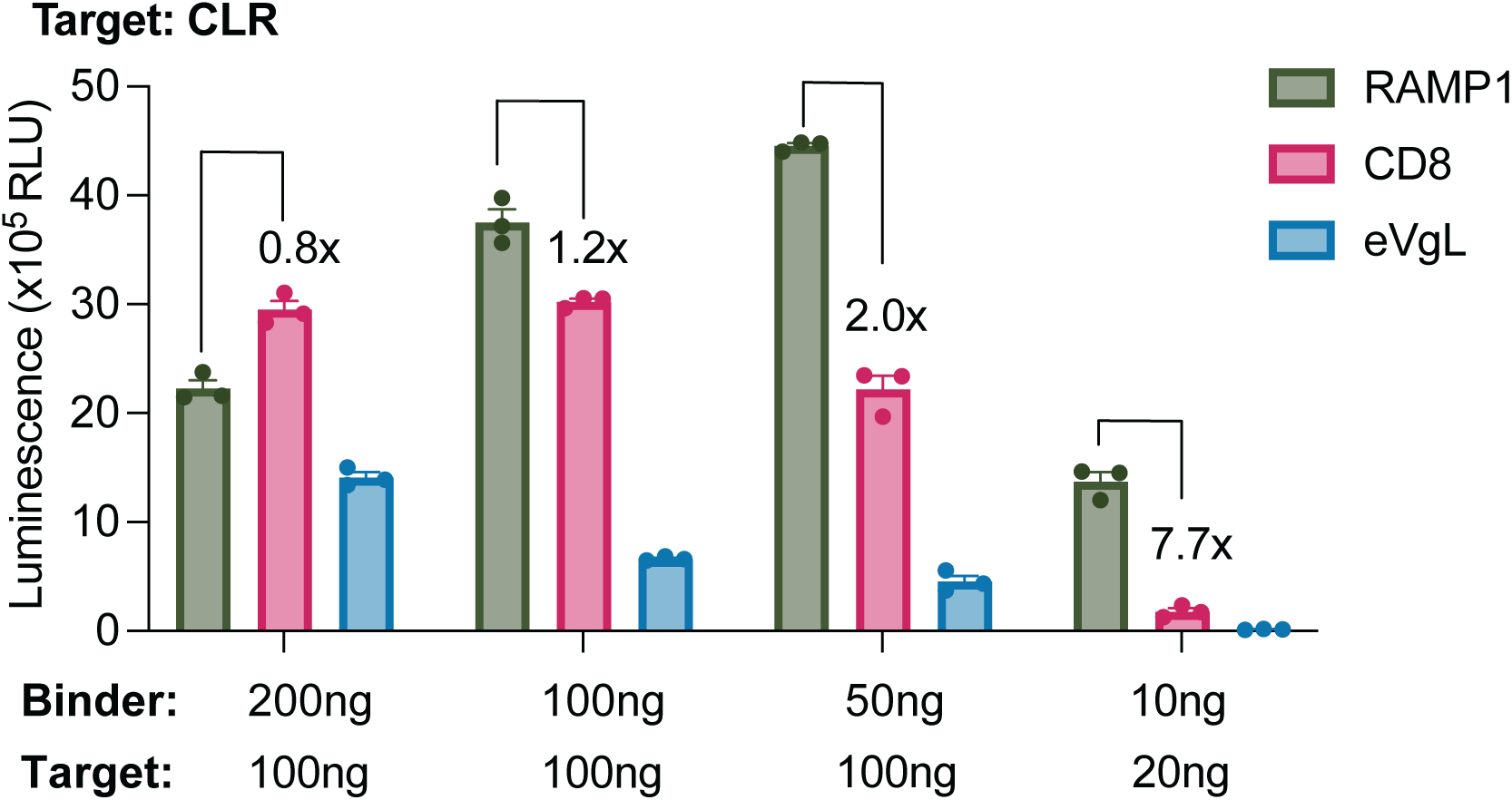
Relationship of membrane protein transfection and dynamic range. CLR-smBit target co-transfected with natural and synthetic TM LgBit-tagged potential binders in two regimes of high and low DNA transfection in HEK293T cells. Left three groups represent 100ng CLR “target” with 200, 100, and 50ng “Binder” RAMP1, and last group represents optimized 20ng CLR + 10ng Binder. Raw Luminescence signals, with fold-change between positive and negative controls noted. N=1 independent experiment. Error bars, SEM.

**Supplementary Figure 3.**
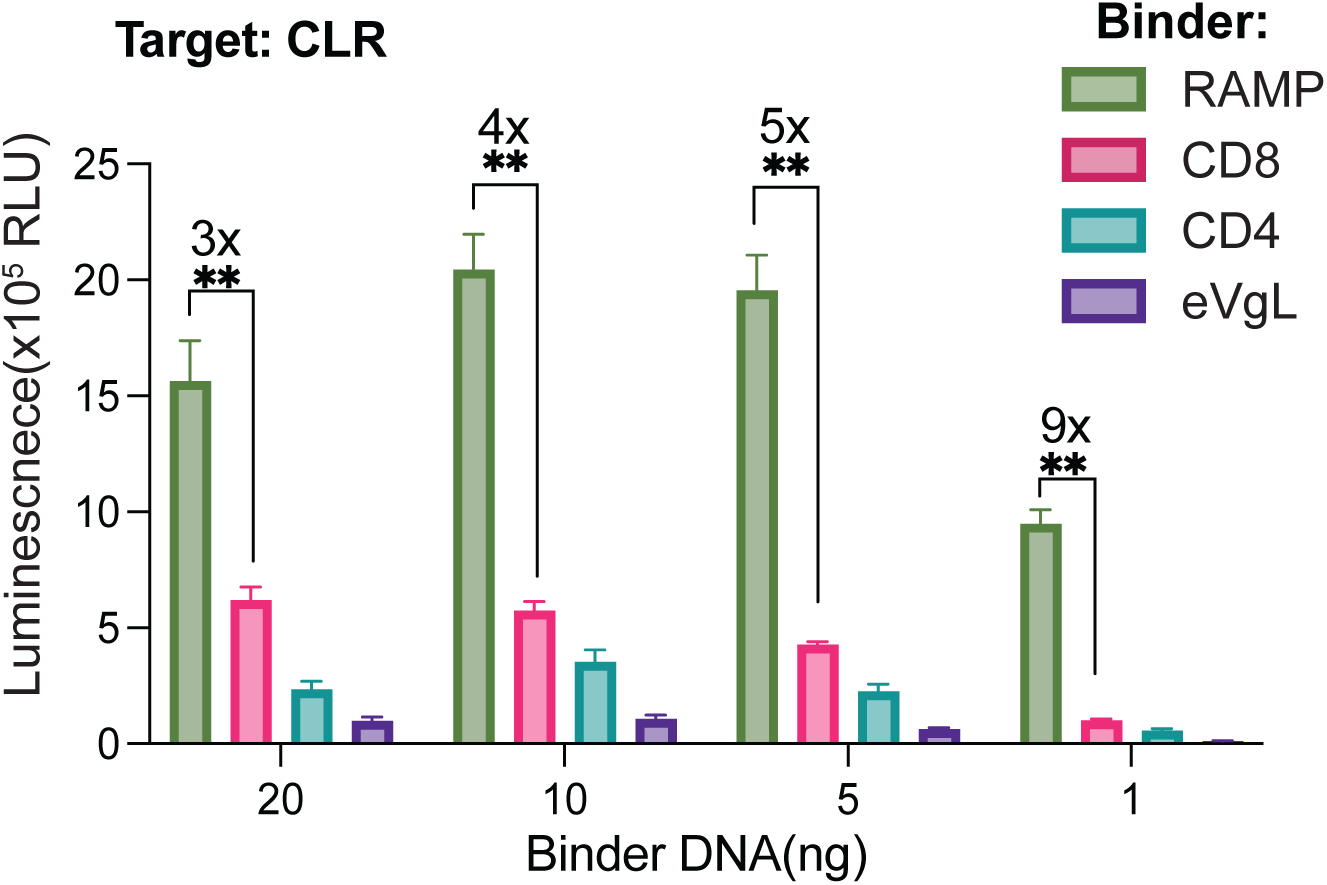
Vehicle DNA does not affect NanoBit dynamic range. NanoBit performed with a variation compared to Figures 1-2 with the “target” CLR-SmBit DNA concentration transfected (10 ng) and titrating “binder” LgBit fusion TM protein constructs (20, 10, 5, 1 ng), by supplementing empty pcDNA as vehicle to have equal final total plasmid DNA amounts in the transfection set at 30 ng total DNA. For example, with 10 ng CLR-SmBit and 1 ng RAMP-LgBit, 19 ng of vehicle pcDNA are added. Luminescence signals with known binder (RAMP1) and non-specific interaction binders (CD8, CD4 and eVgL). Data show n=3 independent biological replicates. Error bars, standard error of the mean. Asterisks indicate P values reaching <0.001 from unpaired t-test comparing signal from CLR-RAMP expressing cells versus that of CLR-CD8a expressing cells.

**Supplementary Figure 4.**
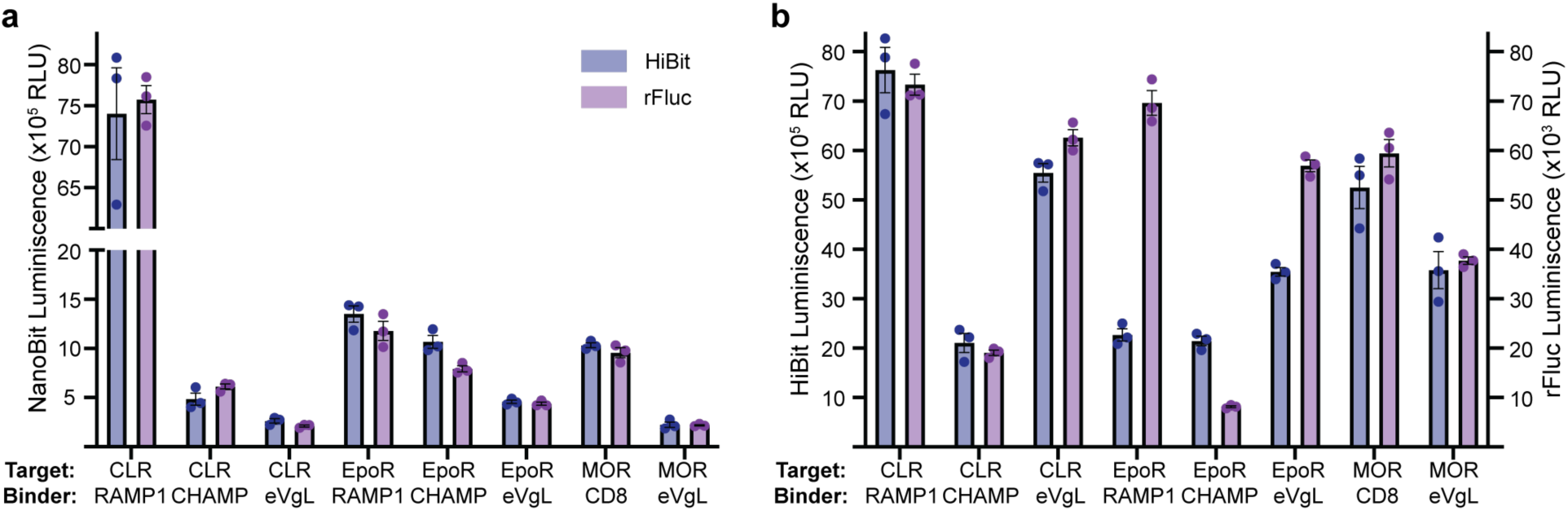
Representative luminescence values of NanoBit, HiBit, and rFluc used for normalization. a. Luminescence values for NanoBit measurements of target-binder pairs. Blue-colored bars and dots indicate LgBit “Binder” expression level normalization by 2-step HiBit measurement. Purple-colored bars and dots indicate LgBit “Binder” expression level normalization by 2-step T2A-rFluc measurement. n=1 trial shown, representative of N=3 independent experiments with 3 technical replicates. Error bars, standard error of the mean. b. Luminescence values indicating LgBit “Binder” expression level as measured by 2-step HiBit (blue) or T2A-rFluc (purple) measured subsequently to the Nanobit PPI measurement. These raw protein expression luminescence values were used to normalized the NanoBit PPI luminescence per sample for the experiment in panel (**a)**.

**Supplementary Table 1.**
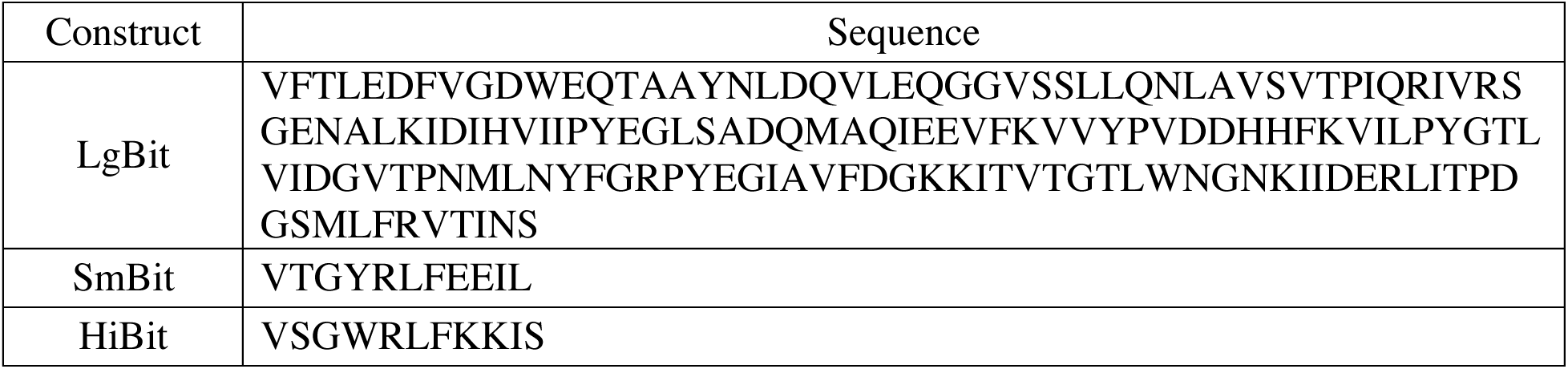
Sequences for LgBit, smBit, HiBit.

**Supplementary Table 2.**
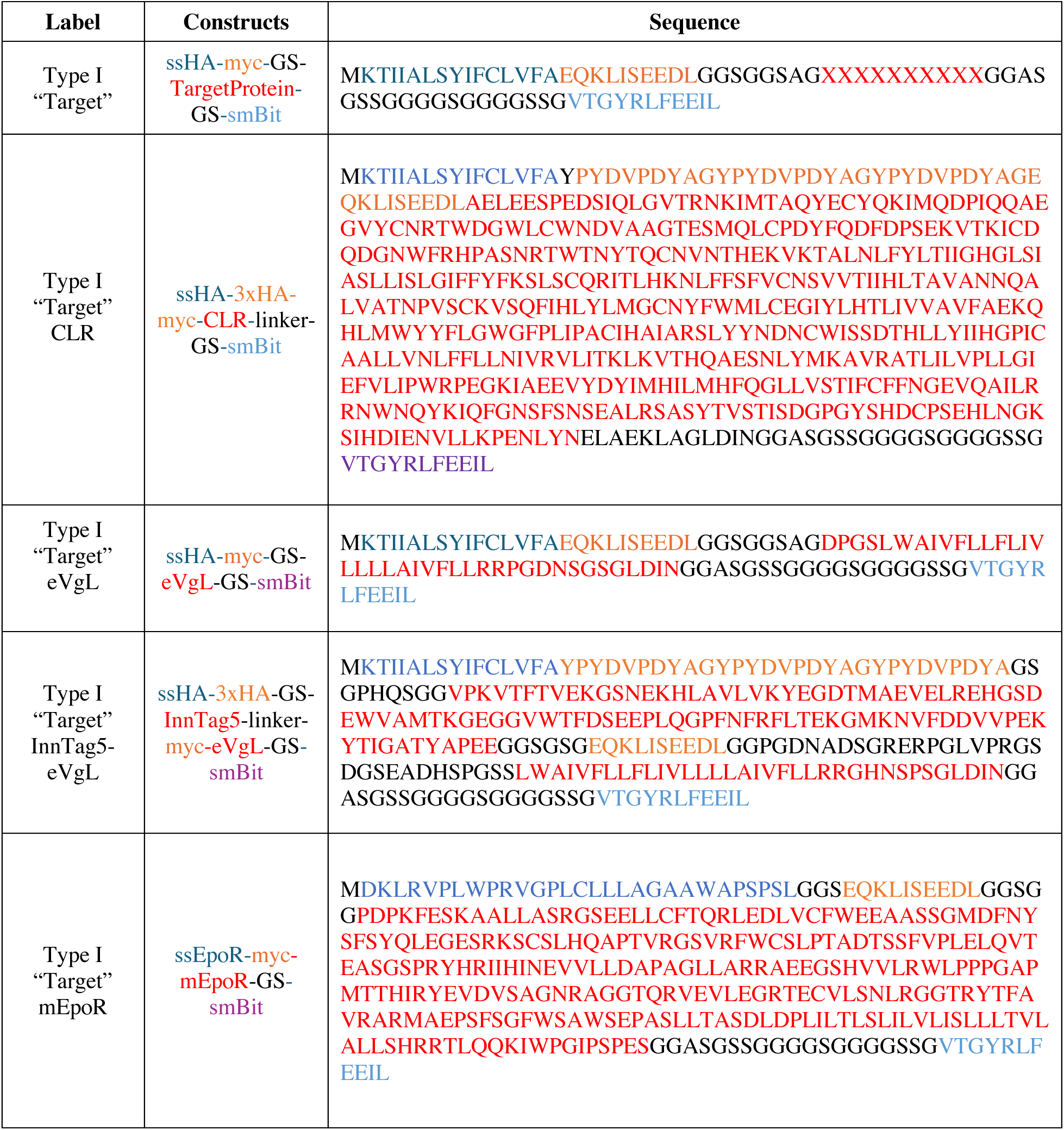
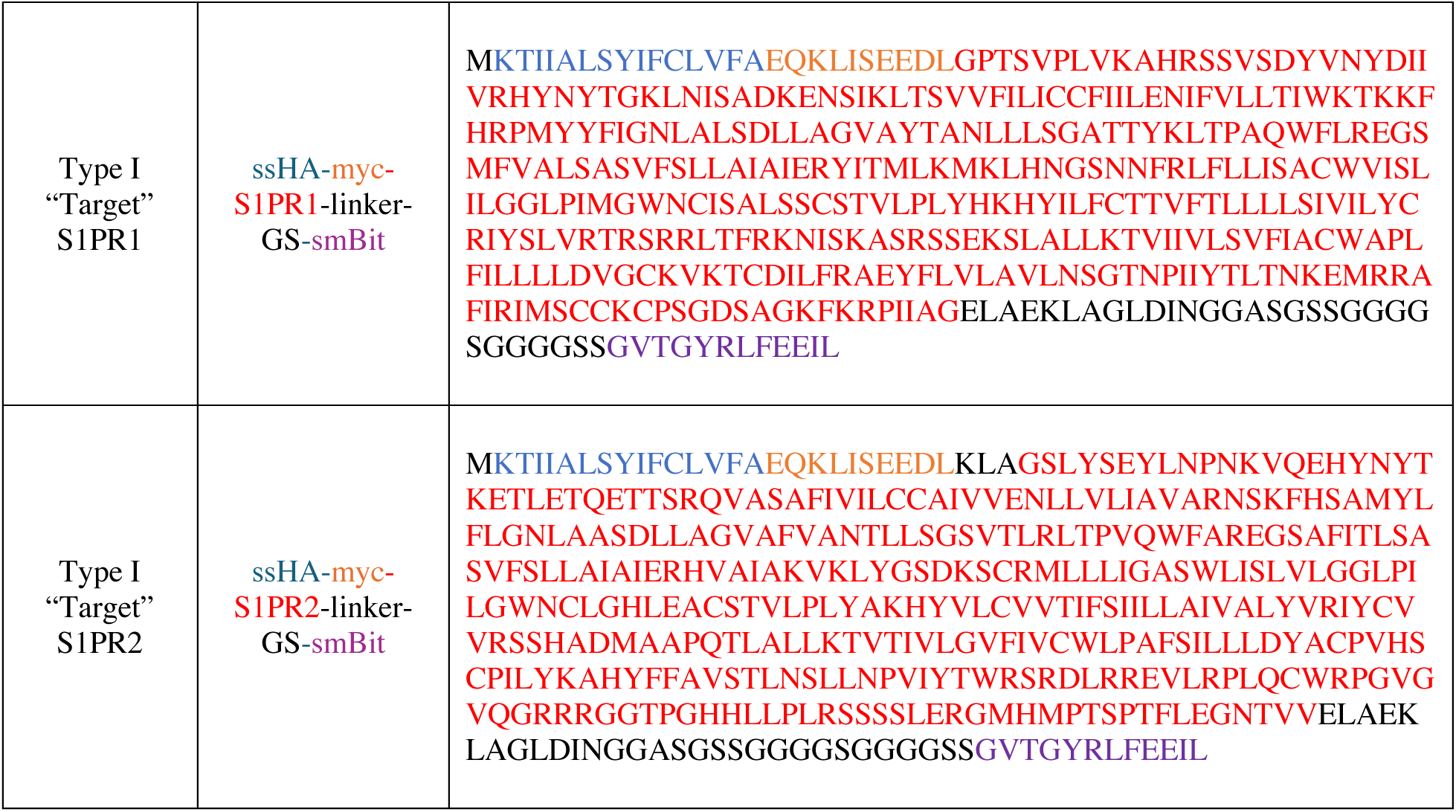
“Target” SmBit-tagged constructs. General expression constructs and representative examples for SmBit-tagged targets utilizing ssHA, HA/myc epitope tag(s), TM protein, and SmBit domain. Examples of eVgL with and without InnTag5 soluble domains are shown. Specific examples of CLR, S1PR1, S1PR2, and mEpoR target constructs are listed.

**Supplementary Table 3.**
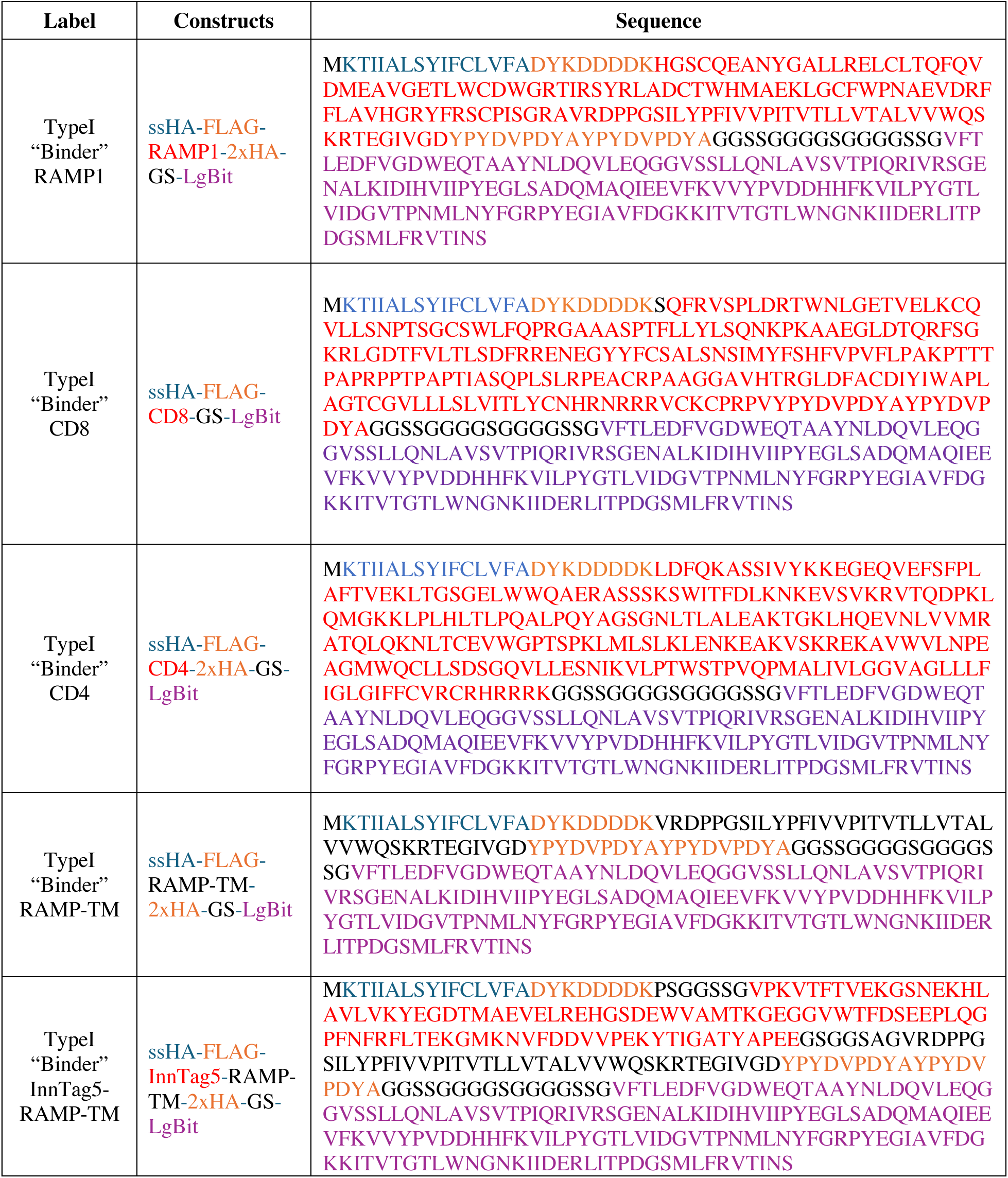
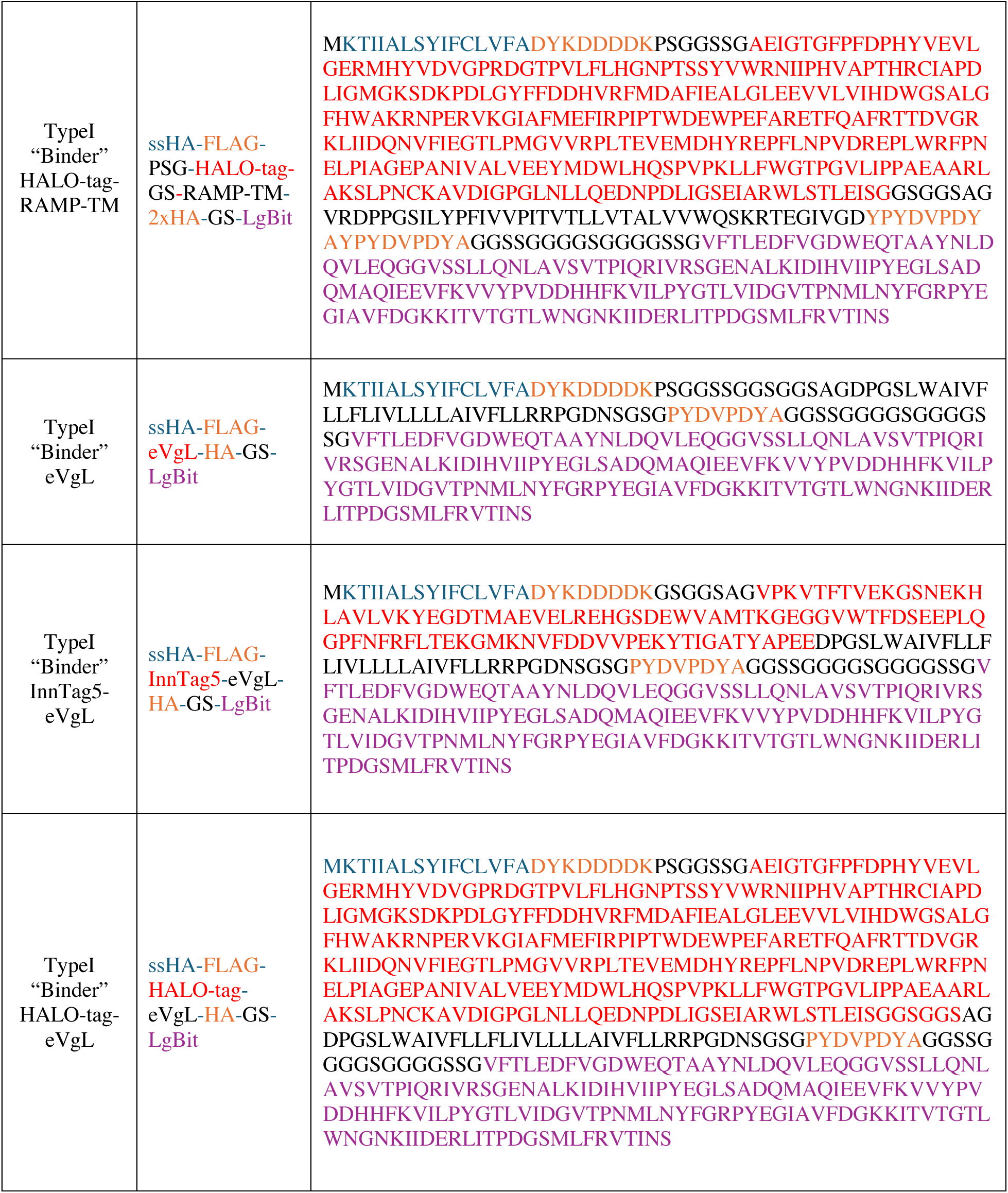
“Binder” LgBit-tagged constructs. Expression constructs for LgBit-tagged targets utilizing ssHA, FLAG epitope tag, TM protein, and LgBit domain. Examples of eVgL and RAMP1 with and without soluble domains (InnTag5 or HALO-tag) are shown. Specific examples of CD8 and CD4 “binder” constructs are listed.

**Supplementary Table 4.**
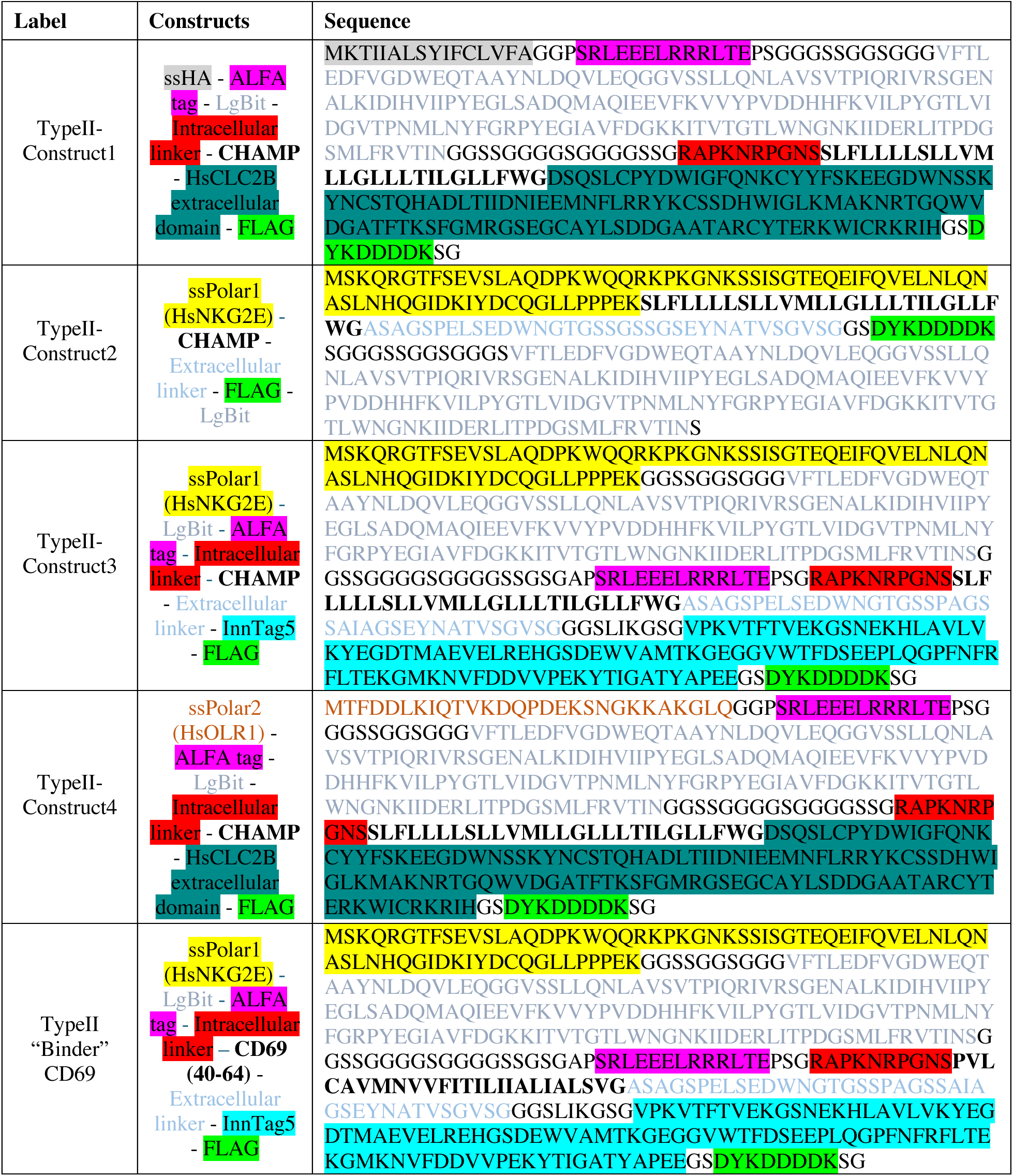
Sequences for Type II constructs.

## References

1 Westerfield, J. M. & Barrera, F. N. Membrane receptor activation mechanisms and transmembrane peptide tools to elucidate them. Journal of Biological Chemistry 295, 1792–1814 (2020).

2 Duart, G., Grau, B., Mingarro, I. & Martinez-Gil, L. Methodological approaches for the analysis of transmembrane domain interactions: A systematic review. Biochim Biophys Acta Biomembr 1863, 183712 (2021). 10.1016/j.bbamem.2021.183712

3 Zoued, A., Duneau, J. P. & Cascales, E. Bacterial One- and Two-Hybrid Assays to Monitor Transmembrane Helix Interactions. Methods Mol Biol 2715, 259–271 (2024). 10.1007/978-1-0716-3445-5_17

4 Dixon, A. S. et al. NanoLuc Complementation Reporter Optimized for Accurate Measurement of Protein Interactions in Cells. ACS Chem Biol 11, 400–408 (2016). 10.1021/acschembio.5b00753

5 Langosch, D., Brosig, B., Kolmar, H. & Fritz, H. J. Dimerisation of the glycophorin A transmembrane segment in membranes probed with the ToxR transcription activator. J Mol Biol 263, 525–530 (1996). 10.1006/jmbi.1996.0595

6 Kolmar, H. et al. Membrane insertion of the bacterial signal transduction protein ToxR and requirements of transcription activation studied by modular replacement of different protein substructures. EMBO J 14, 3895–3904 (1995). 10.1002/j.1460-2075.1995.tb00061.x

7 Armstrong, C. R. & Senes, A. Screening for transmembrane association in divisome proteins using TOXGREEN, a high-throughput variant of the TOXCAT assay. Biochim Biophys Acta 1858, 2573–2583 (2016). 10.1016/j.bbamem.2016.07.008

8 Joce, C., Wiener, A. A. & Yin, H. Multi-Tox: application of the ToxR-transcriptional reporter assay to the study of multi-pass protein transmembrane domain oligomerization. Biochim Biophys Acta 1808, 2948–2953 (2011). 10.1016/j.bbamem.2011.07.008

9 Lindner, E. & Langosch, D. A ToxR-based dominant-negative system to investigate heterotypic transmembrane domain interactions. Proteins 65, 803–807 (2006). 10.1002/prot.21226

10 Su, P. C. & Berger, B. W. A novel assay for assessing juxtamembrane and transmembrane domain interactions important for receptor heterodimerization. J Mol Biol 425, 4652–4658 (2013). 10.1016/j.jmb.2013.07.022

11 Steindorf, D. & Schneider, D. In vivo selection of heterotypically interacting transmembrane helices: Complementary helix surfaces, rather than conserved interaction motifs, drive formation of transmembrane hetero-dimers. Biochim Biophys Acta Biomembr 1859, 245–256 (2017). 10.1016/j.bbamem.2016.11.017

12 Schneider, D. & Engelman, D. M. GALLEX, a measurement of heterologous association of transmembrane helices in a biological membrane. J Biol Chem 278, 3105–3111 (2003). 10.1074/jbc.M206287200

13 Russ, W. P. & Engelman, D. M. TOXCAT: a measure of transmembrane helix association in a biological membrane. Proc Natl Acad Sci U S A 96, 863–868 (1999). 10.1073/pnas.96.3.863

14 Dawson, J. P., Weinger, J. S. & Engelman, D. M. Motifs of serine and threonine can drive association of transmembrane helices. J Mol Biol 316, 799–805 (2002). 10.1006/jmbi.2001.5353

15 Russ, W. P. & Engelman, D. M. The GxxxG motif: a framework for transmembrane helix-helix association. J Mol Biol 296, 911–919 (2000). 10.1006/jmbi.1999.3489

16 Schanzenbach, C., Schmidt, F. C., Breckner, P., Teese, M. G. & Langosch, D. Identifying ionic interactions within a membrane using BLaTM, a genetic tool to measure homo- and heterotypic transmembrane helix-helix interactions. Sci Rep 7, 43476 (2017). 10.1038/srep43476

17 Julius, A., Laur, L., Schanzenbach, C. & Langosch, D. BLaTM 2.0, a Genetic Tool Revealing Preferred Antiparallel Interaction of Transmembrane Helix 4 of the Dual-Topology Protein EmrE. J Mol Biol 429, 1630–1637 (2017). 10.1016/j.jmb.2017.04.003

18 Elazar, A. et al. Mutational scanning reveals the determinants of protein insertion and association energetics in the plasma membrane. Elife 5 (2016). 10.7554/eLife.12125

19 Lis, M. & Blumenthal, K. A modified, dual reporter TOXCAT system for monitoring homodimerization of transmembrane segments of proteins. Biochem Biophys Res Commun 339, 321–324 (2006). 10.1016/j.bbrc.2005.11.022

20 Remy, I. & Michnick, S. W. Clonal selection and in vivo quantitation of protein interactions with protein-fragment complementation assays. Proc Natl Acad Sci U S A 96, 5394–5399 (1999). 10.1073/pnas.96.10.5394

21 Rossi, F., Charlton, C. A. & Blau, H. M. Monitoring protein-protein interactions in intact eukaryotic cells by beta-galactosidase complementation. Proc Natl Acad Sci U S A 94, 8405–8410 (1997). 10.1073/pnas.94.16.8405

22 Paulmurugan, R., Umezawa, Y. & Gambhir, S. S. Noninvasive imaging of protein-protein interactions in living subjects by using reporter protein complementation and reconstitution strategies. Proc Natl Acad Sci U S A 99, 15608–15613 (2002). 10.1073/pnas.242594299

23 Kodama, Y. & Hu, C. D. An improved bimolecular fluorescence complementation assay with a high signal-to-noise ratio. Biotechniques 49, 793–805 (2010). 10.2144/000113519

24 Romei, M. G. & Boxer, S. G. Split Green Fluorescent Proteins: Scope, Limitations, and Outlook. Annu Rev Biophys 48, 19–44 (2019). 10.1146/annurev-biophys-051013-022846

25 Yao, Z. et al. Split Intein-Mediated Protein Ligation for detecting protein-protein interactions and their inhibition. Nat Commun 11, 2440 (2020). 10.1038/s41467-020-16299-1

26 Johnsson, N. & Varshavsky, A. Split ubiquitin as a sensor of protein interactions in vivo. Proc Natl Acad Sci U S A 91, 10340–10344 (1994). 10.1073/pnas.91.22.10340

27 Tebo, A. G. & Gautier, A. A split fluorescent reporter with rapid and reversible complementation. Nat Commun 10, 2822 (2019). 10.1038/s41467-019-10855-0

28 Shekhawat, S. S. & Ghosh, I. Split-protein systems: beyond binary protein-protein interactions. Curr Opin Chem Biol 15, 789–797 (2011). 10.1016/j.cbpa.2011.10.014

29 Cabantous, S., Terwilliger, T. C. & Waldo, G. S. Protein tagging and detection with engineered self-assembling fragments of green fluorescent protein. Nat Biotechnol 23, 102–107 (2005). 10.1038/nbt1044

30 Magliery, T. J. et al. Detecting protein-protein interactions with a green fluorescent protein fragment reassembly trap: scope and mechanism. J Am Chem Soc 127, 146–157 (2005). 10.1021/ja046699g

31 Duart, G. et al. Computational design of BclxL inhibitors that target transmembrane domain interactions. Proc Natl Acad Sci U S A 120, e2219648120 (2023). 10.1073/pnas.2219648120

32 Botta, J., Bibic, L., Killoran, P., McCormick, P. J. & Howell, L. A. Design and development of stapled transmembrane peptides that disrupt the activity of G-protein-coupled receptor oligomers. J Biol Chem 294, 16587–16603 (2019). 10.1074/jbc.RA119.009160

33 Gales, C. et al. Real-time monitoring of receptor and G-protein interactions in living cells. Nat Methods 2, 177–184 (2005). 10.1038/nmeth743

34 Angers, S. et al. Detection of beta 2-adrenergic receptor dimerization in living cells using bioluminescence resonance energy transfer (BRET). Proc Natl Acad Sci U S A 97, 3684–3689 (2000). 10.1073/pnas.97.7.3684

35 Barnea, G. et al. The genetic design of signaling cascades to record receptor activation. Proc Natl Acad Sci U S A 105, 64–69 (2008). 10.1073/pnas.0710487105

36 Kim, M. W. et al. Time-gated detection of protein-protein interactions with transcriptional readout. Elife 6 (2017). 10.7554/eLife.30233

37 Petschnigg, J. et al. The mammalian-membrane two-hybrid assay (MaMTH) for probing membrane-protein interactions in human cells. Nat Methods 11, 585–592 (2014). 10.1038/nmeth.2895

38 Xu, Y., Piston, D. W. & Johnson, C. H. A bioluminescence resonance energy transfer (BRET) system: application to interacting circadian clock proteins. Proc Natl Acad Sci U S A 96, 151–156 (1999). 10.1073/pnas.96.1.151

39 Li, B. et al. High-Throughput NanoBiT-Based Screening for Inhibitors of HIV-1 Vpu and Host BST-2 Protein Interaction. Int J Mol Sci 22 (2021). 10.3390/ijms22179308

40 Peach, C. J., Kilpatrick, L. E., Woolard, J. & Hill, S. J. Use of NanoBiT and NanoBRET to monitor fluorescent VEGF-A binding kinetics to VEGFR2/NRP1 heteromeric complexes in living cells. Br J Pharmacol 178, 2393–2411 (2021). 10.1111/bph.15426

41 Lay, C. S., Kilpatrick, L. E., Craggs, P. D. & Hill, S. J. Use of NanoBiT and NanoBRET to characterise interleukin-23 receptor dimer formation in living cells. Br J Pharmacol 180, 1444–1459 (2023). 10.1111/bph.16018

42 Inoue, A. et al. Illuminating G-Protein-Coupling Selectivity of GPCRs. Cell 177, 1933–1947 e1925 (2019). 10.1016/j.cell.2019.04.044

43 Liang, Y. L. et al. Cryo-EM structure of the active, G(s)-protein complexed, human CGRP receptor. Nature 561, 492–497 (2018). 10.1038/s41586-018-0535-y

44 Chen, H., Qin, Y., Chou, M., Cyster, J. G. & Li, X. Transmembrane protein CD69 acts as an S1PR1 agonist. Elife 12 (2023). 10.7554/eLife.88204

45 Mravic, M. et al. Packing of apolar side chains enables accurate design of highly stable membrane proteins. Science 363, 1418–1423 (2019). 10.1126/science.aav7541

46 Kratochvil, H. T. et al. Transient water wires mediate selective proton transport in designed channel proteins. Nature Chemistry 15, 1012–1021 (2023). 10.1038/s41557-023-01210-4

47 Mravic, M. et al. De novo-designed transmembrane proteins bind and regulate a cytokine receptor. Nature Chemical Biology (2024). 10.1038/s41589-024-01562-z

48 Kotliar, I. B. et al. Multiplexed mapping of the interactome of GPCRs with receptor activity-modifying proteins. Sci Adv 10, eado9959 (2024). 10.1126/sciadv.ado9959

49 Huttlin, E. L. et al. Dual proteome-scale networks reveal cell-specific remodeling of the human interactome. Cell 184, 3022–3040 e3028 (2021). 10.1016/j.cell.2021.04.011

50 Devaraneni, P. K. et al. Stepwise insertion and inversion of a type II signal anchor sequence in the ribosome-Sec61 translocon complex. Cell 146, 134–147 (2011). 10.1016/j.cell.2011.06.004

51 Dou, D., da Silva, D. V., Nordholm, J., Wang, H. & Daniels, R. Type II transmembrane domain hydrophobicity dictates the cotranslational dependence for inversion. Mol Biol Cell 25, 3363–3374 (2014). 10.1091/mbc.E14-04-0874

52 Georgieva, M. V. et al. Inntags: small self-structured epitopes for innocuous protein tagging. Nat Methods 12, 955–958 (2015). 10.1038/nmeth.3556

53 Murase, T. et al. Identification of soluble forms of lectin-like oxidized LDL receptor-1. Arterioscler Thromb Vasc Biol 20, 715–720 (2000). 10.1161/01.atv.20.3.715

54 Tsirigos, K. D., Peters, C., Shu, N., Kall, L. & Elofsson, A. The TOPCONS web server for consensus prediction of membrane protein topology and signal peptides. Nucleic Acids Res 43, W401–407 (2015). 10.1093/nar/gkv485

55 Thumuluri, V., Almagro Armenteros, J. J., Johansen, A. R., Nielsen, H. & Winther, O. DeepLoc 2.0: multi-label subcellular localization prediction using protein language models. Nucleic Acids Res 50, W228–W234 (2022). 10.1093/nar/gkac278

56 Krishna Kumar, K., et al. Negative allosteric modulation of the glucagon receptor by RAMP2. Cell 186, 1465–1477 e1418 (2023). 10.1016/j.cell.2023.02.028

57 Levy, S. & Shoham, T. The tetraspanin web modulates immune-signalling complexes. Nat Rev Immunol 5, 136–148 (2005). 10.1038/nri1548

